# Redefinition of archetypal phytoplankton-associated bacteria taxa based on globally distributed dinoflagellates and diatoms

**DOI:** 10.1101/2023.02.13.528248

**Authors:** Xiaoyu Yang, Guanjing Cai, Runlin Cai, Haifeng Gu, Yuerong Chen, Jianmin Xie, Zhong Hu, Hui Wang

**Author notes:** Corresponding author: Hui Wang. These two authors contributed equally (Xiaoyu Yang, Guanjing Cai).

## Abstract

Bacteria colonizing in the phycosphere formed by phytoplankton exudates play important roles in marine ecosystems, yet their taxonomy is poorly defined. Here, we customized the analytical approaches for the microalga-attached microbiotas from 110 diatom and 86 dinoflagellate samples to reveal key bacterial players and their ecological significance in the phycosphere. The results demonstrated a co-occurrence of host-specificity and conservation of phytoplankton-associated bacterial communities, defined 8 diatom- and 23 dinoflagellate-affiliated characteristic genera, as well as identifying 14 core genera prevalent with phytoplankton populations. Further classification of these 14 core genera into three tiers showed their distinct ecological features regarding occupancy, connectivity and community-stabilizing, whilst also matching their inherent metabolic properties. Our study redefines the archetypal phytoplankton-associated bacteria taxa more specifically up to the genus level, highlighting the significance of rarely noticed bacteria in the phycosphere, which is invaluable when selecting target bacteria for studying phytoplankton-bacteria interactions.

## Introduction

Phytoplankton are primary aquatic producers forming the basis of most marine food webs and are critical to the stability of marine ecosystems^1^. These organisms can fix 30 - 50 billion metric tons of carbon per year, equating to about 40% of the total fixed carbon making them important regulators of global carbon^2^. Besides the biomass of phytoplankton, these fixed carbons are also transformed into extracellular products released to a certain area outside the cells, which forms a diffusive layer defined as the “algal sphere” now known as “phycosphere” ^3, 4^. Phycosphere with high concentration of molecules attracts heterotrophic bacteria, providing a critical interface for the occurrence of phytoplankton-bacteria interaction based on the exchange of nutrients and metabolites. Generally, exudates and lysates of phytoplankton cells serve as nutrient sources or metabolic precursors for bacteria^5,6,7^, while bacteria provide vitamins, ammonium salts, and other trace elements to support the growth of phytoplankton^8, 9^. In addition, bacteria can kill algal cells by synthesizing specific metabolites^10^ or inhibit growth of microalgae by competing for nutrients^11^. These diverse interactions of mutualism, commensalism, competition and antagonism^1^ deeply regulates growth of both populations. Since heterotrophic bacteria can consume more than 50% of the primary productivity fixed by phytoplankton, phytoplankton-bacteria interaction have been acknowledged as one of the most fundamental ecological relationships in the aquatic environment which vastly influences the major biogeochemical cycles on Earth^12^.

By exploring the framework of substance exchange for phytoplankton-bacteria interaction, the metabolic bond between bacteria and phytoplankton explains why the composition of bacterial communities can be shaped by phytoplankton species and algal bloom phases^4^. However, global surveys reveal a relatively conserved group called “archetypal phytoplankton-associated bacterial taxa” that live in symbiosis with phytoplankton, mainly including members of the classes Alphaproteobacteria, Flavobacteriia and Gammaproteobacteria^13–15^. For example, amplicon data from two different species of *Pseudo-nitzschia* indicated species specificity between bacteria and diatoms, but members of *Rosebacter*, Gamma-proteobacteria and Flavobacteria coexisted with both diatom species persistently^16^; Molecular investigations of bacterial diversity in six diatom cultures revealed different bacterial phylotypes associated with different diatom genera, while Alphaproteobacteria, Bacteroidetes and small amount of β-proteobacteria were always the most prominent in the phycosphere of all diatoms^17^. The existence of these conserved bacterial groups suggests that there are some universal metabolic mechanisms regulating the phytoplankton-bacteria interactions. However, current studies on phytoplankton-associated bacteria communities contain obvious shortcomings that limit our cognition. First, *in situ* studies relying on physical grading can neither separate influences of different phytoplankton species nor precisely isolate metabolic communications between bacteria and phytoplankton with both contact and non-contact dependent interactions. Therefore, the contamination of unrelated phytoplankton and bacteria cannot be eliminated, hampering data accuracy. Second, although purposeful isolation and long-termed cultivation of phytoplankton strains can avoid interference of miscellaneous phytoplankton species and selectively enrich phytoplankton-associated bacterial communities, most studies were limited to focusing only on a few genera in specific phylum since obtaining phytoplankton colonies remain difficult. Lack of data results in the inability to define a unified rule that applies to bacteria communities in diverse phycosphere. As such, current studies can only define “archetypal phytoplankton-associated bacterial taxa” at a higher taxonomic level such as class. Limited data regarding these conserved groups on more precise taxonomic levels implies that there is yet to be a reputable reference on phylogenetic-metabolic correlation of high validity to guide research on molecular mechanisms. It is, necessary to precisely identify the conserved “core” microbiota that is tightly associated with broadly representative phytoplankton, to support in-depth studies on bacteria-phytoplankton interactions, with ecological significance.

To tackle these limitations, 196 globally distributed phytoplankton colonies affiliated to photosynthetic Dinophyceae (Dinoflagellates) and Bacillariophyta (Diatoms) were collected; both of which are representative phylum with global distribution and account for more than half of the total biomass of phytoplankton in the ocean^18^. The 16S rRNA gene amplicons of bacterial communities in the phycosphere were sequenced and analyzed based on multiple ecologically relevant statistics to identify the core genera that conservatively presented with dinoflagellates and diatoms and showed prominent ecological significance. The results, verified by the data of *in situ* bloom samples, provided strong evidence for the absolute dominance of specific bacteria, thereby creating a solid foundation for phytoplankton-bacteria interactions to be studied further via either cellar or global perspectives.

## Results

### Overall composition of bacterial communities

A total of 10,476,100 reads from 196 samples were clustered into 3,593 Amplicon Sequence Variants (ASVs). Among them, 2,307 ASVs were from dinoflagellate samples (*n*=86) while 1,194 ASVs were from diatom samples (*n*=110). The manual annotation with the EzBioCloud Database (EZ) was chosen for further analysis since it provided more extensive species information than that with the SILVA database (Supplementary Fig. 1). The taxa with abundance greater than 1% of the total sequences were summarized and visualized at each taxonomic level (Fig. 1c). Coinciding with the current understanding of “archetypal phytoplankton-associated bacterial taxa”, our samples were also dominated by three major classes: Alphaproteobacteria, Gammaproteobacteria and Flavobacteriia. Each major class was mainly composed of two leading bacterial clades, resulting in the six top families being Roseobacteraceae, Flavobacteriaceae, Crocinitomicaceae, Rhodobacteraceae, Alteromonadaceae and Oceanospirillaceae. These families contributed >70% of the total sequence reads. Most of these families contained only one or two genera, such as *Neptuniibacter* in Oceanospirillaceae, while Roseobacteraceae and Flavobacteriaceae were highly diverse, each consisting of at least five highly abundant genera. Among them, *Marivita* (10.4%) and *Shimia* (3.7%) accounted for nearly half of Roseobacteraceae, while *Winogradskyella* (3%) and *Polaribacter* (2.6%) were the most abundant genera in Flavobacteriaceae.

**Figure 1.**
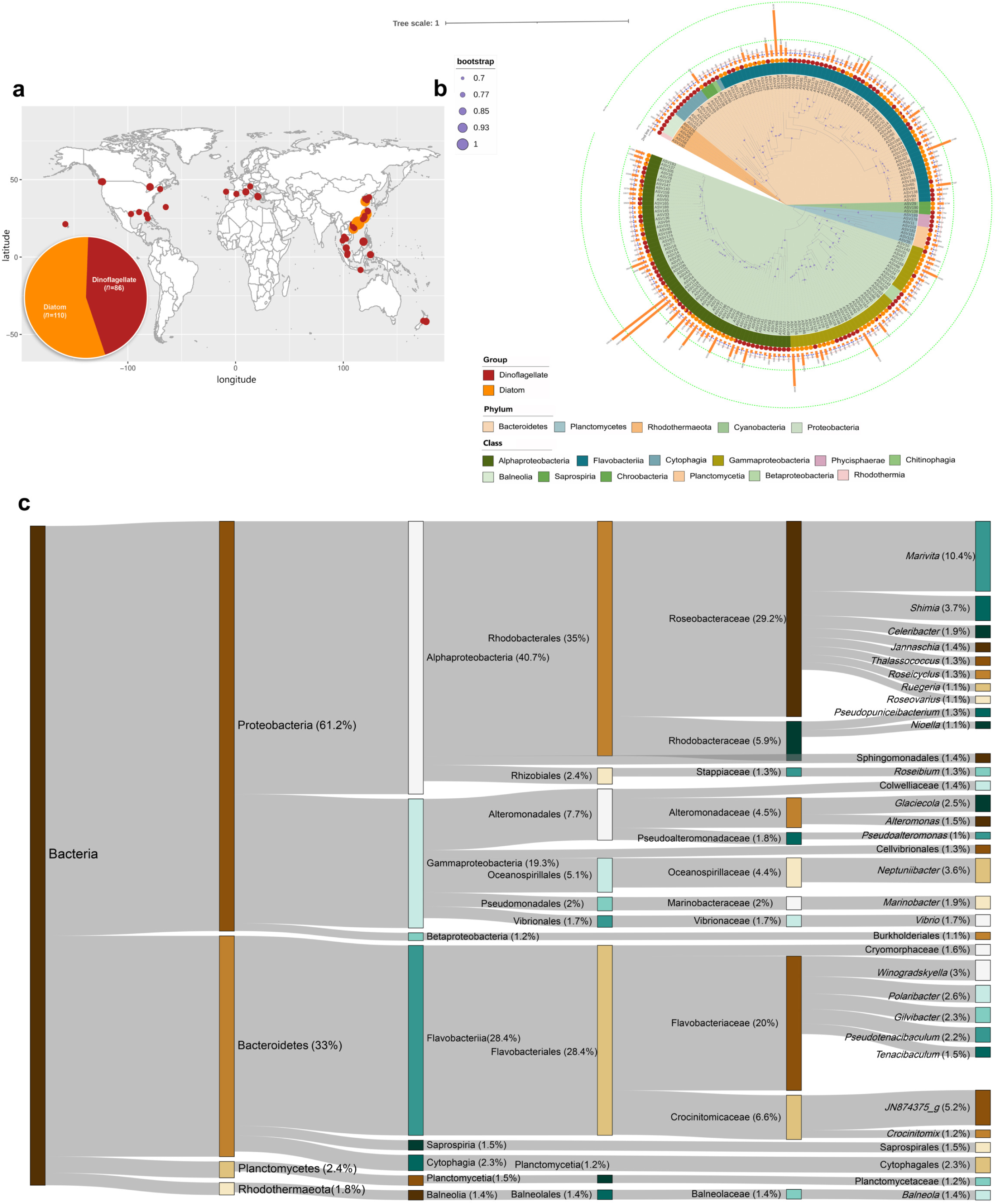
Overview of bacterial community composition in the physcophere. **a,** Sample Information. The sampling locations are shown by dots on the map, with orange-red indicating dinoflagellate samples and orange-yellow indicating diatom samples, and the size of the dots indicating the number of samples. **b,** A phylogenetic evolutionary tree of 200 high-abundance ASVs constructed based on the maximum likelihood method. The background color of the evolutionary branch indicates the phylum, the innermost circle indicates the Class, the pie chart in the middle circle indicates the distribution proportion of ASVs in the dinoflagellate and diatom groups, and the bar chart in the outermost circle indicates the sequencing amount of ASVs in all samples. **c,** Composition of bacterial taxa at each taxonomic level and the proportion in total sequencing volume.

Furthermore, digging into the origin and taxonomic information of the 200 high-abundance ASVs (>0.1%) (Fig. 1b), 79 of them were distributed in the phycosphere of both dinoflagellate and diatom, such as the most sequenced ASVs - ASV54 and ASV134, both affiliated to *Marivita*. Similarly, ASVs with prominent abundance, including ASV66 (*Shimia*), ASV117 (*Nioella*), ASV167 (*Alteromonas*), ASV114 (*Winogradskyella*) and ASV197 (*Roseovarius*) also homogeneously distributed, thereby indicating the conservatism of dominant taxa across different phycosphere. However, some high-abundance ASVs showed a strong preference for either dinoflagellate or diatom, as evidenced by ASV71 (*Neptuniibacter*), ASV44 (*Glaciecola*), ASV42 (*Polaribacter*) and ASV1 (*Celeribacter*) that existed with diatoms only, as well as ASV34 (*Gilvibacter*), ASV13 (*Balneola*), ASV88 (*Psychrosphaera*) and ASV163 (*Methylophaga*) that existed with dinoflagellates only. Considering that these diatom- or dinoflagellate-specific bacteria showed outstanding average abundances which were not even diluted by the samples from unpreferred phytoplankton during calculation, host phytoplankton could drive the assembly of bacterial community in the phycosphere.

### Differences of bacterial communities in the phycosphere of dinoflagellates and diatoms

To clarify influences of host phytoplankton on bacterial communities, genus statistics, α-, β-diversities, and co-occurrence networks of bacterial assemblages grouped by dinoflagellates and diatoms were analyzed and compared first. Summarizing the genera that formed the bacterial communities, there were only 182 genera shared by both groups, accounting for 29.8% of the 610 genera in the dinoflagellate group and 70.3% of the 259 genera in the diatom group (Supplementary Fig. 2a). The existence of significantly more dinoflagellate-specific genera (428) compared with diatom-specific genera (77) also suggested higher community diversity in the former group, which could be confirmed by the significantly higher richness and evenness indices of α-diversity than those of the diatom group (Fig. 2a). However, the co-occurrence networks showed that bacterial ASVs in the diatom group had more connections than those in the dinoflagellate group. Diatom-associated bacteria networks contained 106 nodes and 258 edges, while dinoflagellate-associated bacteria networks only contained 76 and 207 respectively (Fig. 2c, d, Supplementary Data3). Members in the network of each group differed significantly; only five genera were shared by both groups. In addition, highly connective nodes varied greatly in different networks. For example, nodes belonging to *Glaciecola* and *Neptuniibacter* with great negative correlation to other nodes in the diatom group and ASVs belonging to *Hoeflea* and *JF514260_g* with universal positive correlation to other nodes in the dinoflagellate group only appeared in their respective groups. These results strongly suggested that different host phytoplankton recruited different bacterial population, which was verified by the β-diversity distance matrix based on the Unweight-Unifrac distance of phylogenetic relationship which showed two well-clustered groups with only partial intersection (Fig. 2b). Permutational multivariate analysis of variance (PERMANOVA) analysis confirmed that the between-group differences (R^2^ =0.129, F=28.447, Pr(>F) =0.001) were significantly greater than the within-group differences, indicating the decisive role of host phytoplankton in shaping bacterial community.

**Figure 2.**
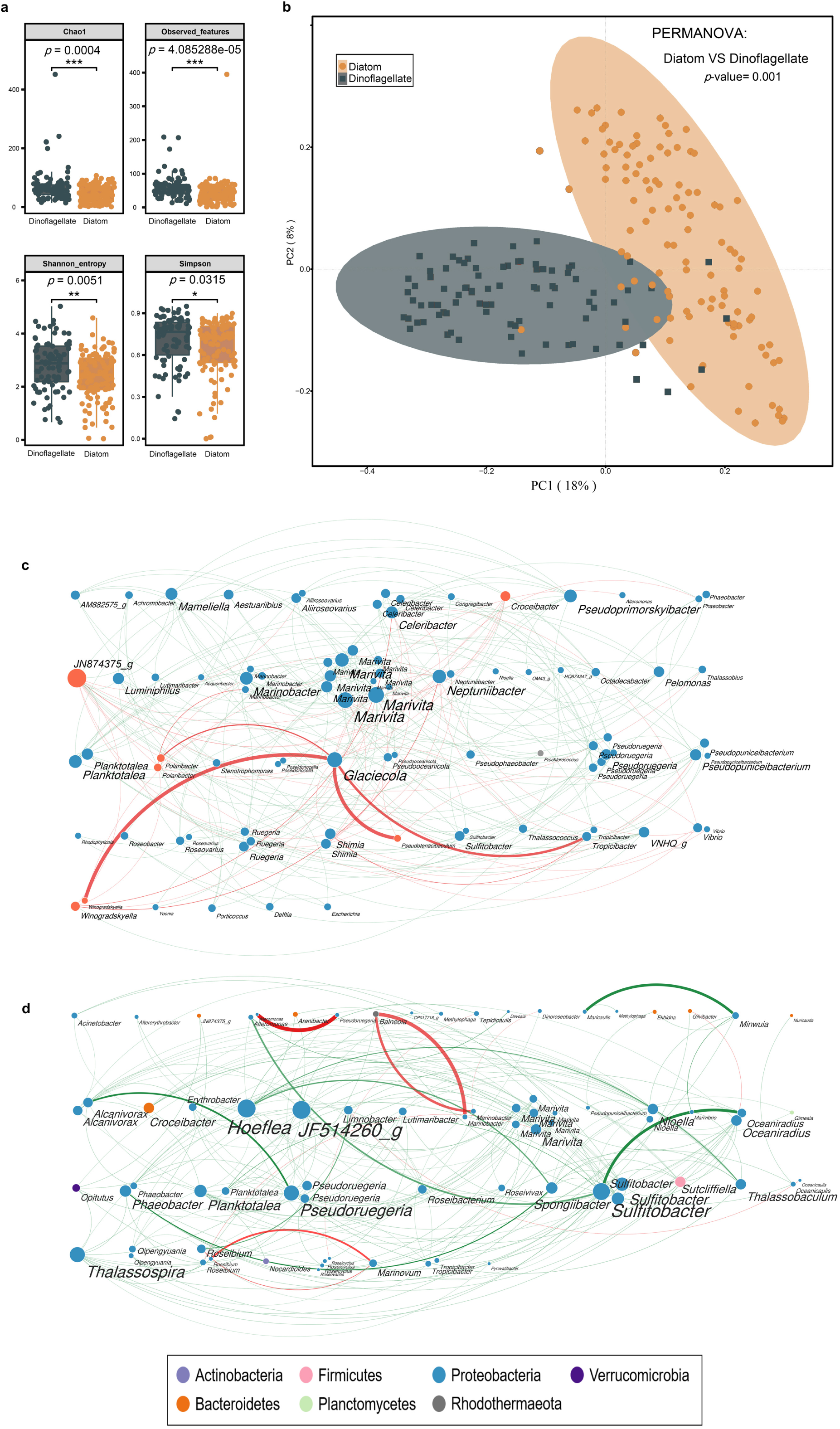
Differences of bacterial communities in the physcophere of dinoflagellates and diatoms. **a,** Alpha diversity, grey boxes represent the four diversity indices respectively, and the *p* values of significance of differences between the two groups were tested by Wilcoxon rank-sum test. **b,** Beta diversity. color and shape represent grouping, method using Unwei_Unifrac distance matrix, PERMANOVA for significance of differences test (*P* = 0.001). **c** and **d,** Ecological network constructed by ASVs of diatom group (c) and dinoflagellate group (d). ASVs are clustered at the genus level, the size of the solid circles indicates the degree of the nodes; the color of the nodes indicates the classification level of the phyla; the thickness of the edges indicates the size of the weights; the color of the edges indicates the different interrelationships between the ASVs: green indicates copresence, red indicates mutual exclusion.

Linear discriminant analysis of effect sizes (LEfSe) and random forest (RF) model were then combined to identify characteristic genera responsible for general differences of bacterial community between the two groups. 86 genera with linear discriminant analysis (LDA) scores greater than 3 were identified (*p*<0.05), of which 30 were distinctly more abundant in the diatom group and 56 in the dinoflagellate group (Supplementary Data 4). Five trials of tenfold cross-validation of RF model were conducted to clarify the optimal bacterial biomarkers of 50 genera with top MeanDecreaseGini (MDG) values, according to the mean of minimum cross-validation error (Supplementary Fig. 3). Taking the intersection of the results of LEfSe and RF, 32 genera showed strong preferences for the diatom and the dinoflagellate groups respectively (Fig. 3a-c). The affiliations of these genera were further verified by the receiver operating characteristic (ROC) curves (Fig. 3d, e). Except for *Ruegeria* for the dinoflagellate group, the remaining genera all showed >50% area under curve (AUC) even with consideration of the entire confidence interval (95%). Based on these analysis, 8 diatom- and 23 dinoflagellate-affiliated characteristic genera were finally defined. The AUC values also indicated that *Polaribacter* had the strongest preference for the diatom (71.4%), while *Roseibium* was the most “loyal” dinoflagellate-affiliated genus (90.5%). It should be noted that 7/8 of the diatom- and 3/8 of dinoflagellate-affiliated characteristic genera, had mean abundances higher than 1% in corresponding groups, such as *Neptuniibacter* accounted for 6.5% in the diatom group and *Gilvibacter* accounted for 5.2% in the dinoflagellate group. These highly abundant characteristic genera might have closer interactions with their corresponding hosts.

**Figure 3.**
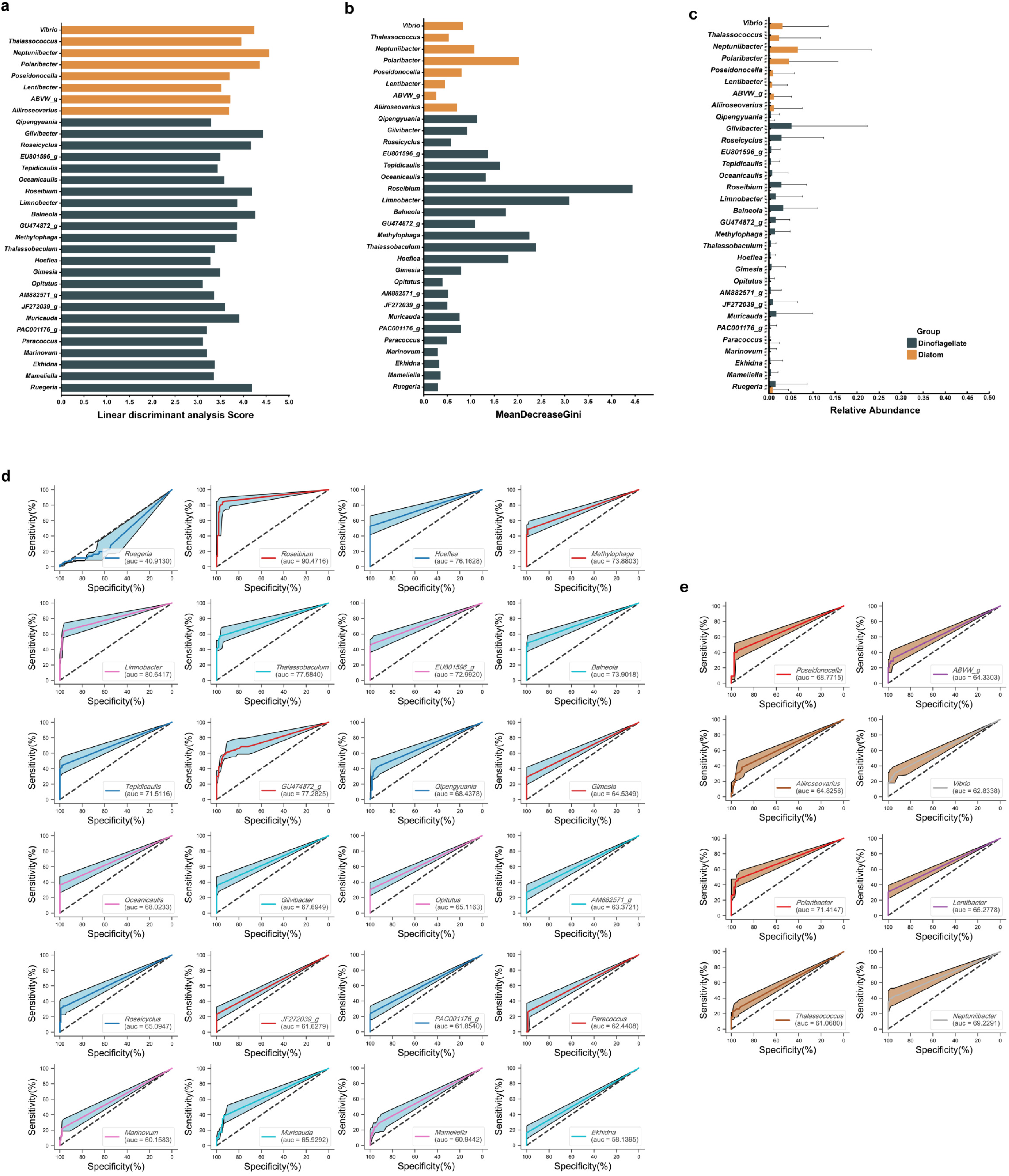
Characteristic genera in the phycosphere of dinoflagellates and diatoms. Linear Discriminant Analysis (LDA) scores (a) and MeanDecreaseGini (b) for the 32 characterized genera, and their relative abundance in each sample of the dinoflagellate and diatom groups(c), Wilcoxon Rank Sum test for significance of differences. Colors represent different groups and stars represent significance levels, *** *P* ≤ 0.001, ** *P* ≤ 0.01, * *P* ≤ 0.05, no star and transparent shading indicates *P* > 0.05 (not significant). **d, e** Receiver operating characteristic curve (ROC curve) for the characteristic genera of dinoflagellate (d) and diatom (e) group. The area under the ROC curve (AUC) greater than 0.5 proved that the genus has some diagnostic value for different host microalgae. Meanwhile, the closer the area under the ROC curve is to 1, the better the authenticity of the diagnostic test. The background color part indicates 95% confidence interval.

### Identification of the core genera across various phycosphere

It should be noted that differences between diatom- and dinoflagellate-associated bacteria communities were statistically significant, but not huge: the 482 dinoflagellate-specific genera accounted for only 11.3% of total abundance of dinoflagellate-associated bacteria, while abundance of the 77 diatom-specific genera was even lower (7.0%) (Supplementary Fig. 2b), indicating low occupancies of phytoplankton-specific bacteria; Shannon and Simpson indices of bacterial communities in the dinoflagellate group were only 14.9% and 8.6% higher than those in the diatom group, exhibiting minor difference on community diversity (Fig. 2a); key parameters of the co-occurrence networks of diatom and dinoflagellate-associated bacteria showed similarity (Fig. 2c, d, Supplementary Data3), such as the mean degree (4.868 and 5.447, respectively) and clustering coefficient (0.331 and 0.337, respectively), demonstrating similar topologies of the networks. The minor differences illustrate that the diatom- and dinoflagellate-associated bacteria communities could share certain similarities and suggest that some conserved bacteria would robustly exist in the phycosphere regardless of host phytoplankton. The conservation of phytoplankton-associated bacterial communities lay the foundations for identifying the core bacteria.

In addition to being conservative in the phycosphere, core bacteria should also be ecologically impactive. Therefore, the first step to reveal core genera was to find ecologically impactive genera (EIGs) with at least one of the three specialties: high abundance, high frequency, and high bio-connectivity. To avoid biases caused by the differentiation effects of host phytoplankton on bacterial communities, EIGs in the diatom and dinoflagellate groups were identified separately. There were 27 high-abundance genera (HAGs) in the dinoflagellate group and 27 HAGs in the diatom group (Fig. 4a, b). Eight HAGs were shared by the two groups, including *Marivita*, *JN874375_g*, *Shimia*, *Pseudotenacibaculum*, *Alteromonas*, *Winogradskyella*, *Marinobacter*, and *Tenacibaculum*, while remaining HAGs differed between groups. In comparison, high-frequency genera (HFGs) were much fewer with more than half of them overlapping with HAGs in the corresponding groups. In the dinoflagellate group, there were 15 HFGs, 10 of which were also designated as HAGs, such as *Roseibium* with the highest frequency of 86% and a high abundance; ranking 6th. The HFG count in the diatom group dropped to only four, while two of them, *Marivita* and *Marinobacter*, were also HAGs in both diatom and dinoflagellate groups. Similarly, 9 of the 22 high bio-connectivity genera (HCGs) in the dinoflagellate group and 14 of the 24 HCGs in the diatom group were also HAGs or HFGs in the corresponding groups (Supplementary Data3). Nonetheless, additional analysis on HCGs included some typical phytoplankton-associated genera, such as *Phaeobacter*, *Thalassospira* in the dinoflagellate group and *Ruegeria* in the diatom group. In summary, merging the HAGs, HFGs and HCGs in corresponding groups, 40 and 47 EIGs were identified in the diatom and dinoflagellate group, respectively.

**Figure 4.**
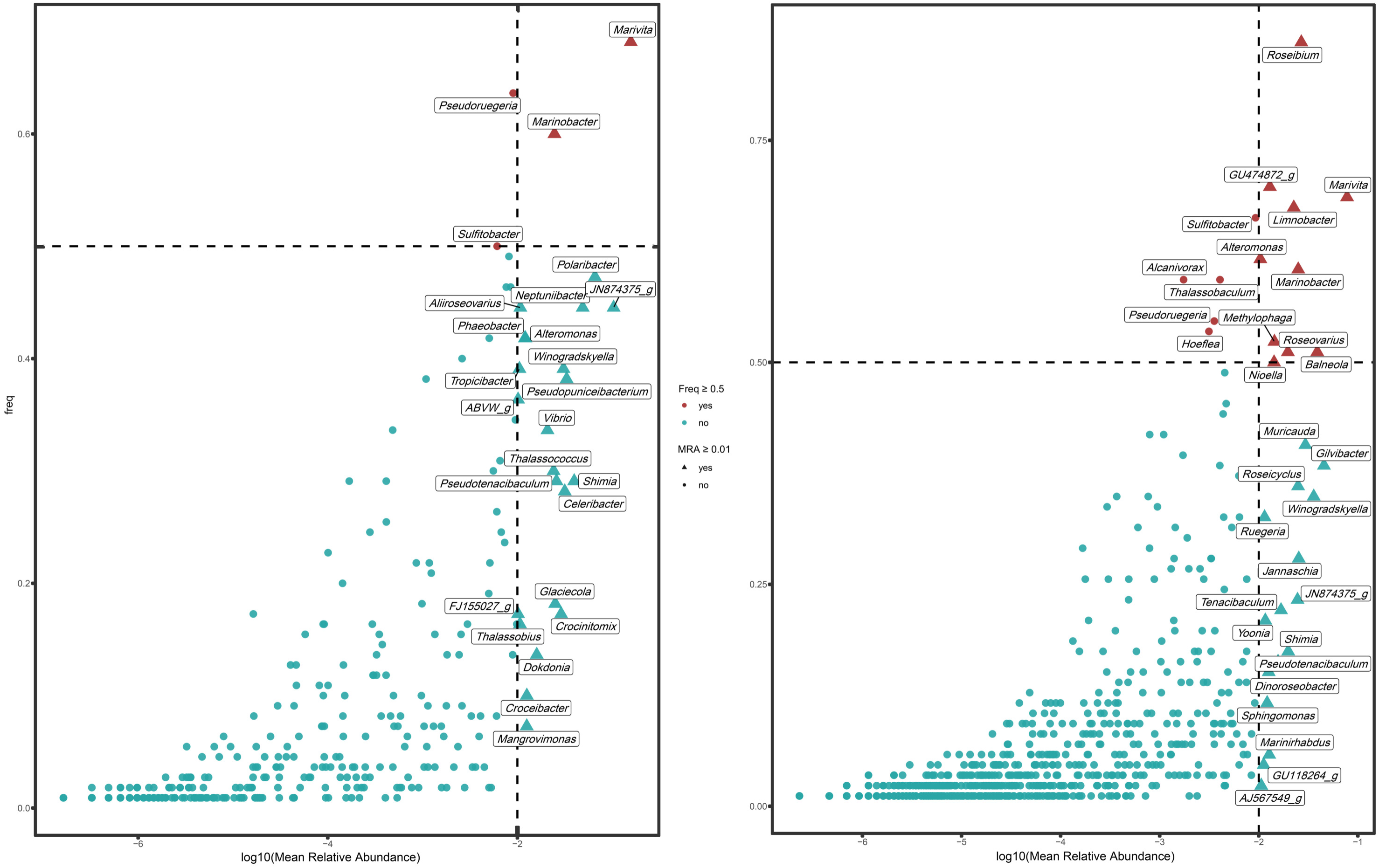
Mean relative abundancies and distribution frequencies of genera in the diatom (a) and dinoflagellate (b) groups. The triangular dots indicate genera with the mean relative abundances >1% (HAGs), and the red dots indicate genera with the distribution frequencies >50% (HFGs).

Taking the intersection of EIGs of the two groups, 14 bacterial genera (Table1) were designated as the core genera that could have universal ecological impact across various phycosphere. Statistics showed that 90% of 196 samples contained at least two core genera (Supplementary Fig. 4a), confirming their prevalence in the phycosphere. These core genera were further categorized into four Types (Table1), according to how their specialties presented in each phytoplankton group. *Marivita* and *Marinobacter* were classified as Type-A, as they possessed more than two specialties in each phytoplankton group. *Pseudoruegeria* and *Sulfitobacter* were both HFGs and HCGs in both phytoplankton groups and therefore classified as Type-B. Type-C consisted of eight genera, including *JN874375_g*, *Alteromonas*, *Winogradskyella*, *Roseovarius*, *Shimia*, *Croceibacter*, *Ruegeria*, and *Pseudotenacibaculum*, which all showed two specialties in one group but only one specialty in the other group. Interestingly, *Ruegeria* had distinct specialties in different phytoplankton groups: it was designated as HFG and HCG in the diatom group but HAG in the dinoflagellate group. *Tenacibaculum* was HAG only and *Planktotalea* was HCG only in both groups, which made them Type-D. The existence of these four Types suggested that different core genera may play different roles in influencing bacterial communities.

**Table 1.**
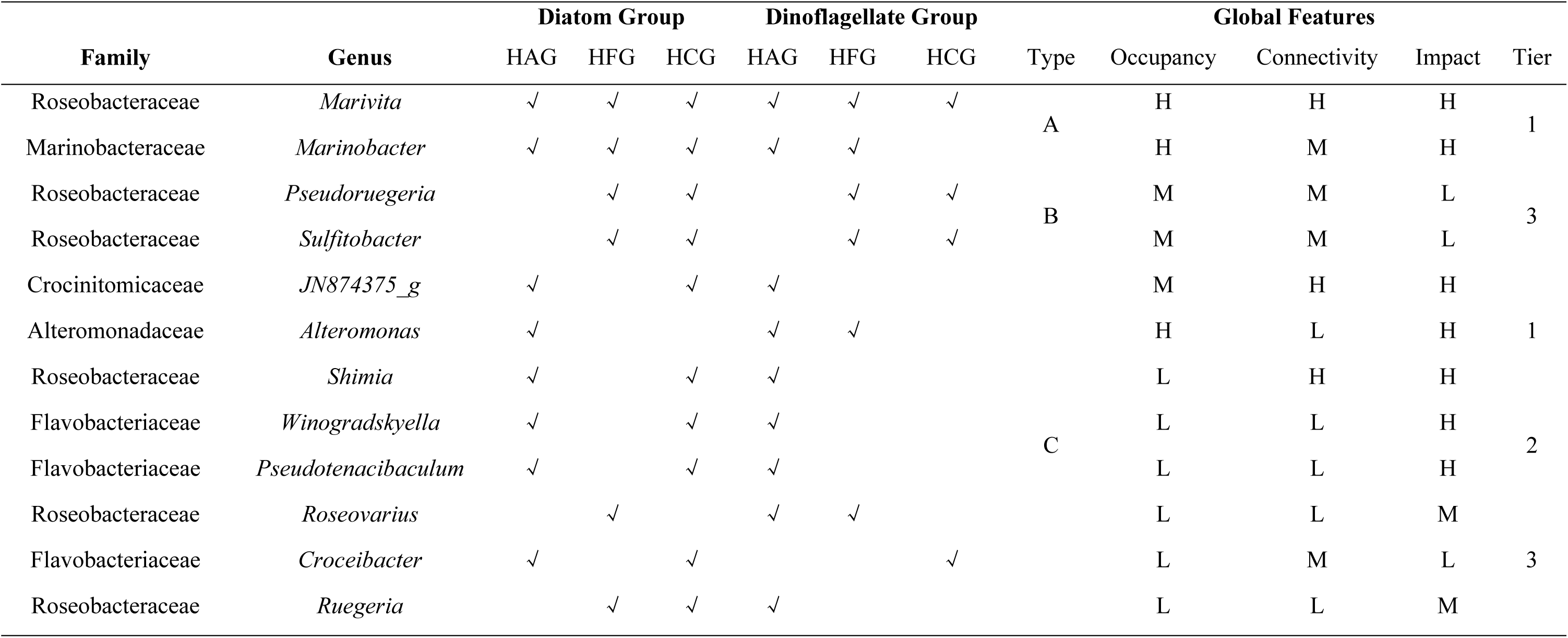
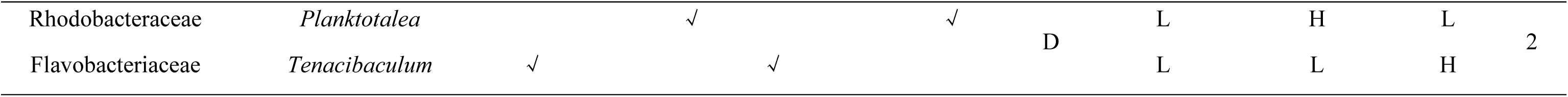
The 14 core genera in the phycosphere bacterial community of dinoflagellates and diatoms. The presence of “√” indicates that the genus satisfies the corresponding characteristic. “H”, “M” and “L” represent the high, medium and low attributes of the global features respectively. Marivita, Marinobacter, JN874375_g, Alteromonas and Shimia, which had at least two “high features”, were classified as Tier-1; Tenacibaculum, Pseudotenacibaculum, Planktotalea and Winogradskyella, which had only one “high feature”, were classified as Tier-2; Sulfitobacter, Roseovarius, *Croceibacter*, *Ruegeria* and *Pseudoruegeria*, which had no “high feature”, were classified as Tier-3.

### Classification of the core genera and their ecological niches

Features exhibited by the subset-based analysis on each phytoplankton group could not demonstrate the universal ecological impact of core genera in a broader range of phycosphere. Therefore, comprehensive analysis based on the complete set of all 196 samples were carried out. Ubiquity-abundance curves, global co-occurrence network and attribution of Bray-Curtis similarity were analyzed to evaluate the global features of each core genus from three aspects: occupancy in bacterial community, connectivity to other genera and impact to community stabilization (Fig. 5, Supplementary Data3). By mapping distributions of core genera for the specified three aspects, Types of core genera showed some correlation to global feature levels – however, some interesting inconsistencies were also noted (Table1). Members of Type-A were both high-occupancy and impact. *Marinobacter* also showed medium-connectivity, while *Marivita* was the most connective, showing highly positive correlation with various non-core genera. The results confirmed the strong dominance of Type-A genera. Members of Type-B were both medium-connectivity and occupancy but had almost no impact to community stabilization (less than 1% contribution to Bray-Curtis similarity), suggesting that these two high ubiquities but low abundance genera might affect only a small range of closely related bacteria. Although members of Type-D were both low-occupancy, each of them exhibited one salient feature of either high-impact (*Tenacibaculum*) or high-connectivity (*Planktotalea*), indicating their strong expertise on influencing bacterial community structure. In contrast, the features of Type-C members were much more complicated. *JN874375_g* showed two “high features” (connectivity and impact) and one “medium feature” (occupancy), just like the Type-A members. *Alteromonas* and *Shimia* also possessed two “high features”, but showed low connectivity and occupancy, respectively. *Winogradskyella* and *Pseudotenacibaculum* that only had one “high feature” were similar with the Type-D members, joining *Tenacibaculum* as high-impact genera. *Roseovarius*, *Croceibacter*, and *Ruegeria* only got two “low features” and one “medium feature” (impact, connectivity and impact, respectively), resulting in those without any “high feature” like the Type-B members. These results illustrated that genera showing more specialties in individual phytoplankton groups (such as Type-B and partial Type-C members) did not necessarily contribute to a more prominent role in a broader range of phycosphere. Instead, genera with specific expertise (such as Type-D members and most Type-C members) might contribute greater to the adaptation of bacterial community to different phytoplankton hosts. According to how many “high features” the core genera possessed, they were rearranged as three Tiers (Table1).

**Figure 5.**
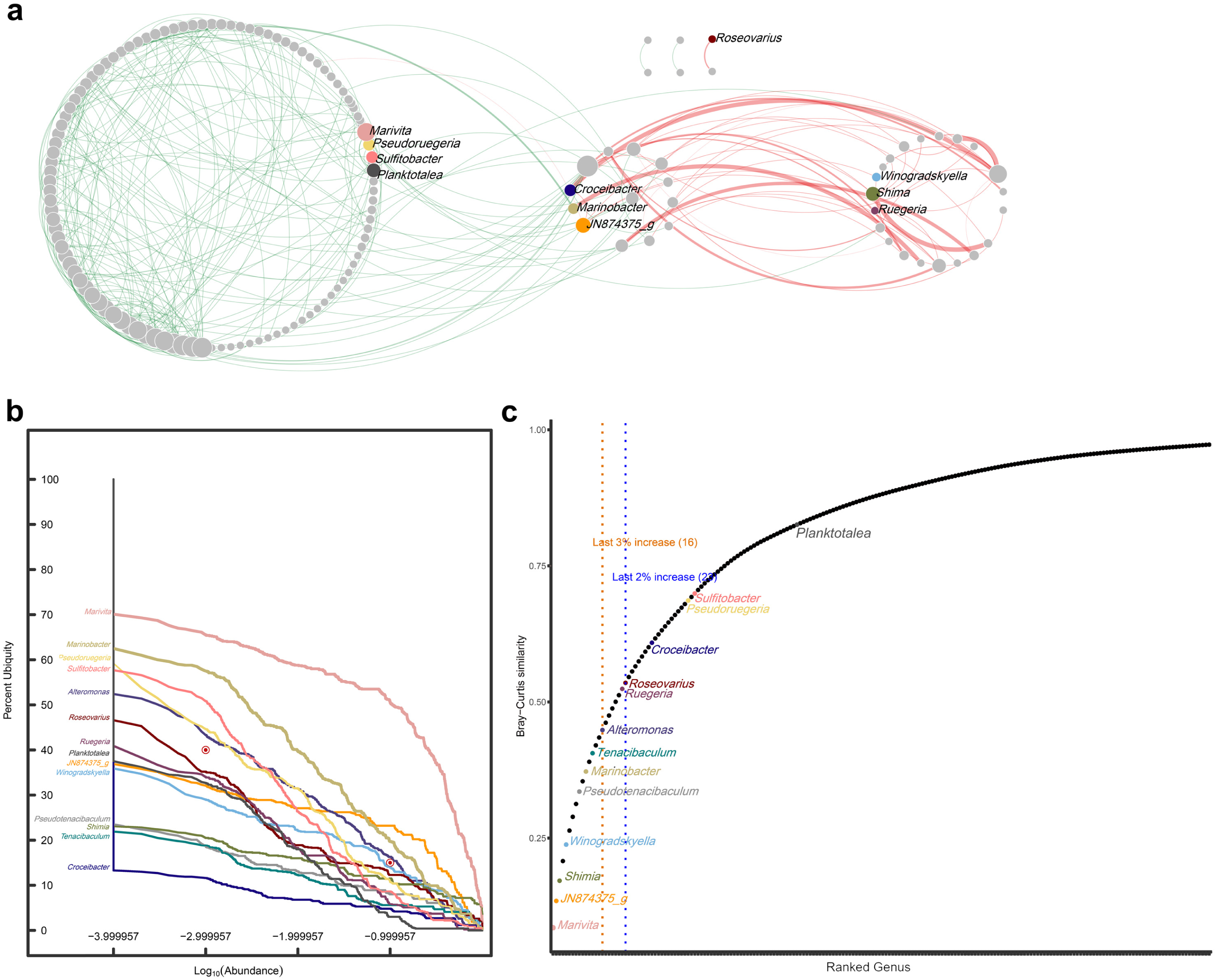
Global features of 14 core genera. **a,** Ecological network of all genera of the microalga-attached (MA) sample. Solid dots represent genera and the size of the dots indicates the degree of nodes; 14 core genera are represented in color, non-core genera are represented in gray; the color of the edges indicates the interaction between genera; the nodes are clustered into three categories according to the interaction relationship: the left circle indicates the interaction are co-occurring; the middle circle indicates the interaction are co-occurring and mutually exclusive; the right circle indicates the interaction are only mutually exclusive. **b,** Core genera Ub-Ab Plot. The Ub-Ab plot for the 14 core genera was constructed by multiplying the ubiquities together for each of core genera along all the abundances. two pairs of abundance versus ubiquity cutoffs (0.01% | 40% and 1% | 15%) were used to determine the genera with low abundance but high ubiquity, and those with high abundance but low ubiquity. **c,** Bray-Curtis similarity for the whole dataset. The ordering of each genus is presented according to the “Method”. The orange and blue line distinguishes core taxa by the last 3% and 2% increase in explanatory value by Bray-Curtis similarity. 14 core genera are represented in color, non-core genera are represented in gray.

Further analysis showed that the core microbiota consisting of the 14 core genera had a distinct ecological niche compared with the non-core microbiota. First, *Levins’* index showed no significant difference of ecological niche width among the 14 core genera (*P* = 0.125, one-way ANOVA), and overall niche width of core microbiota was significantly higher than that of non-core microbiota (*P* =4.5e-10, Wilcox test) (Fig. 6a), indicating higher generality of core microbiota on utilizing resources. The ecological niche overlap index also indicates that competition is more intense within core genera than with non-core genera (Fig. 6b). In most cases, only 5-9 core genera existed simultaneously, suggesting inevitable competition among core genera and potential functional redundancy of core microbiota (Supplementary Fig. 4a). Interestingly, the niche overlaps of Tier-1 vs. 3 and Tier-2 vs. 3 were greater than that of Tier-1 vs. 2 (Fig. 6c), which suggested that the “lower features” of Tier-3 may be attributed to severe competition within core microbiota. Finally, due to the great geographical isolation among the samples, the iCAMP analysis (Fig. 6d) showed that dispersal limitation (73.8%) was the main force causing the differentiation of bacterial communities in the phycosphere, while homogenous selection by host phytoplankton (18.1%) was responsible for the convergence. Compared to non-core microbiota (12.7%), core microbiota was significantly more homogenously selected (23.7%, *P* <0.001, t-test), confirming its close interaction with host phytoplankton. Interestingly, both Tier-1 (18.8%) and Tier-3 (13.1%) showed higher homogenous selection than non-core microbiota, while Tier-2 was much less influenced by selection (8.0%) and dispersal (7.5%) and highly driven by ecological drift (84.5%), demonstrating its random fluctuation in abundance and relatively weak connection with host phytoplankton. The potential differences on the connection with phytoplankton amongst the Tiers also suggests different positioning of ecological niches, which might be determined by their respective phytoplankton-associated metabolic capabilities.

**Figure 6.**
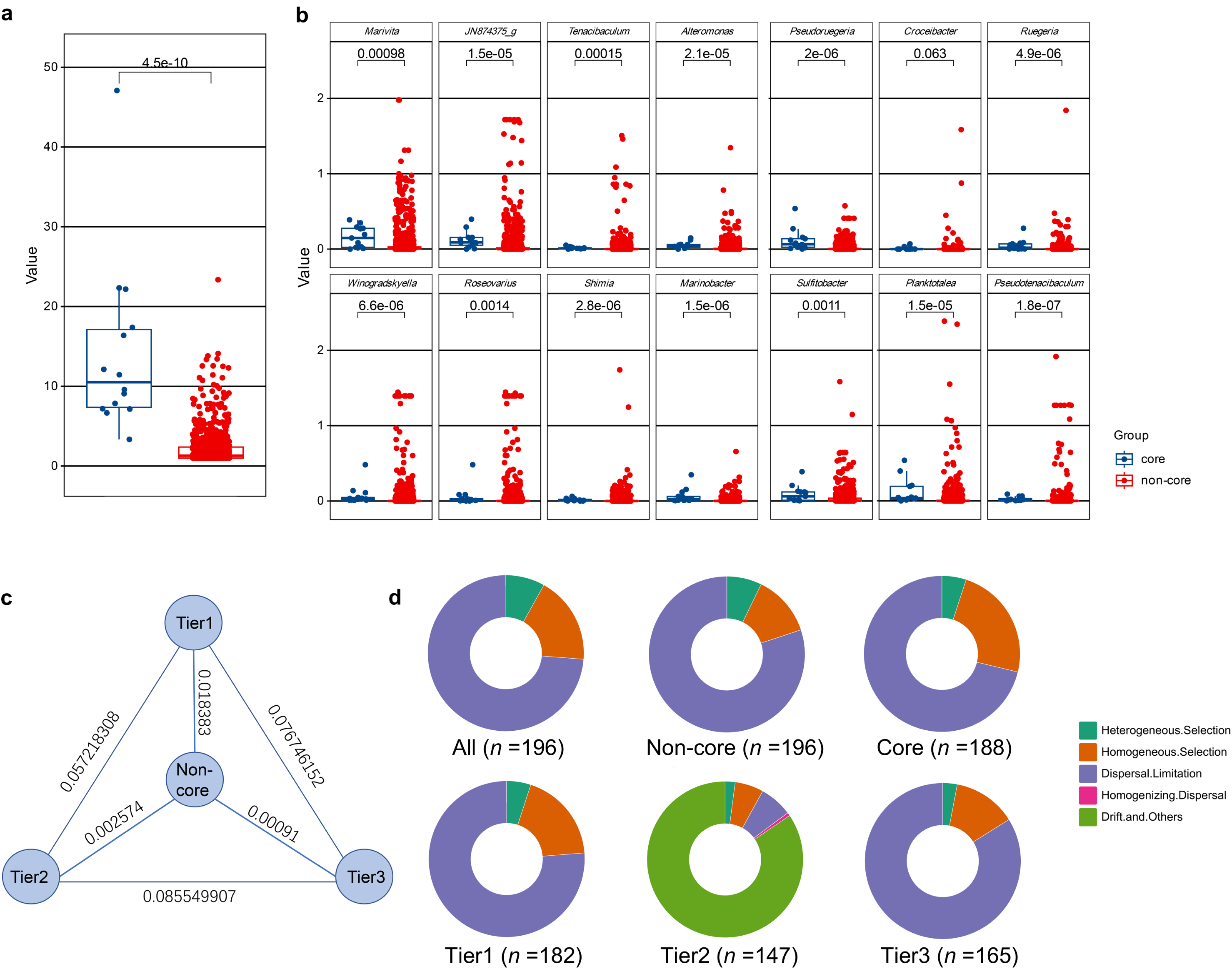
Ecological niches of core and non-core genera. **a,** Ecological niche widths of core and non-core genera based on ‘Levins’ niche width index. **b,** Overlap index of ecological niches of each core genus with other core and non-core genera. Significant differences were determined by Wilcoxon Rank Sum Test, *p* value is displayed as a numeric value. **c,** Overlap of ecological niches between any two of the three Tier groups and non-core genus group. **d,** Ecological processes of community assembling based on different subgroups of microbiotas. Colors distinguish 5 different ecological processes; the size of the solid ring indicates the proportion of each ecological process.

### Verification of the core genera in the *in situ* algal bloom

To verify the ecological significance of the core genera, the *in situ* bloom data from ‘Kabeltonne’ were reanalyzed following the same procedures. A total of 54,419,508 reads from the 137 particle-attached (PA) samples were clustered into 43,554 ASVs belonging to 3,819 genera, and a total of 10,090,181 reads from the 91 free-living (FL) samples were clustered into 7,170 ASVs belonging to 1,968 genera. Since bacteria in the PA samples were more likely to be phytoplankton-associated, they were compared with bacteria in the microalgae-attached (MA) samples (this study) to reveal the roles of the core genera. The results demonstrated that the PA and MA samples shared 687 genera (Fig. 7a), 134 of which had a mean relative abundance >0.1% (Fig. 7d). All 14 core genera belonged to the 134 high abundant genera and each PA sample contained at least five core genera (Supplementary Fig. 4b). In addition, 11 of the 14 core genera existed in more than 50% of the *in situ* samples, especially *JN874375_g*, *Tenacibaculum*, *Winogradskyella*, *Roseovarius*, *Marinobacter*, *Sulfitobacter* that had a high incidence of over 90%, indicating the robust presence of core microbiota in the bloom area. Each core genus also showed higher mean relative abundance in the PA samples than in the FL samples, suggesting potential dispersal of core microbiota from phycosphere to surrounding waters (Fig. 7d, Supplementary Data 5). Core microbiota also had wider niche width than non-core microbiota in the PA samples. The gap was greater than that shown in the MA samples (Fig. 7b), which indicated the prominent resource utilization capability of core microbiota in the bloom area. Similarly, although the difference was minor, core microbiota in the bloom area was still significantly more affected by homogenous selection than non-core microbiota (29.13% vs. 28.61%, *P* =0.022, paired t-test) (Fig. 7c). The consistent characteristics in the PA and MA samples highlighted the conservation and ecological significance of the core microbiota in the phycosphere.

**Figure 7.**
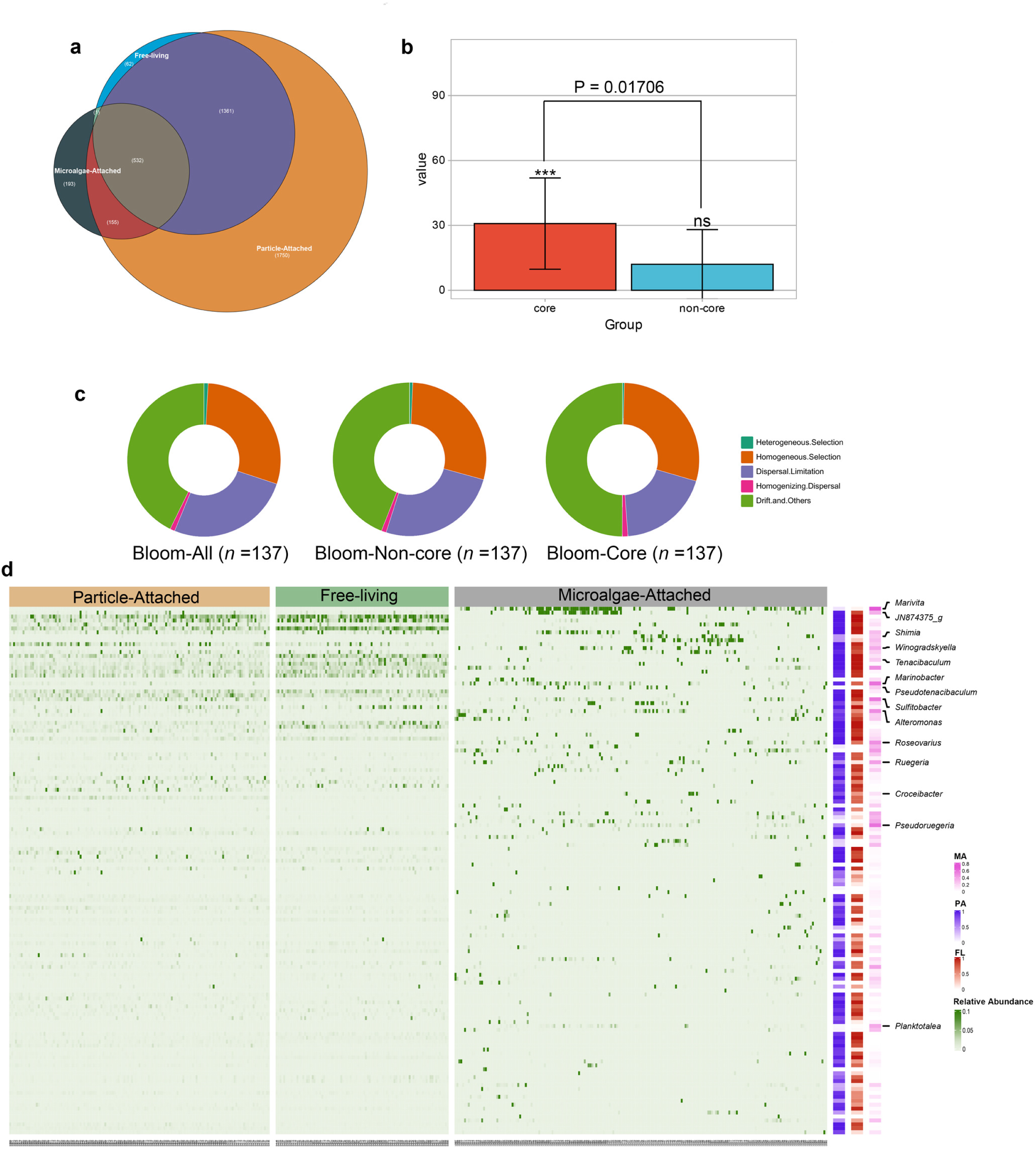
Distribution of core genera in *in situ* bloom data. **a,** Venn diagram showing the numbers of genera in the particle-attached, free-living and microalgae-attached group. **b,** The niche widths of core and non-core genera in the *in situ* bloom samples. **c,** Ecological processes of community assembling based on different subgroups of microbiotas in the *in situ* bloom samples. Colors distinguish 5 different ecological processes; the size of the solid ring indicates the proportion of each ecological process. **d,** Relative abundance of 134 genera in 424 samples. The three-column heatmap (right) indicates the frequency of distribution of each genus in three groups, with light to dark colors indicating low to high frequencies.

## Discussion

Current understanding on “archetypal phytoplankton-associated bacterial taxa” has stagnated at the acknowledgement of the three major groups on vague class level for years^13–15^. Considering the complexity of the phycosphere, summarizing the universal rules to classify bacterial populations may prove challenging. One of the fundamental questions is which has a greater impact on bacterial communities in the phycosphere, the interspecific interaction or environmental condition? Here we performed principal coordinate analysis (PCoA) analysis on the samples from East China Sea (ES), Yellow Sea (YS) and South China Sea (SS) (Supplementary Fig. 5), where both diatoms and dinoflagellates were isolated, to evaluate the influence of environmental condition, using the geographical location of sampling as a surrogate. Although PERMANOVA showed a *p* value <0.001, the three highly overlapped groups and the low R^2^ (0.05428) indicated that geographical differences could barely explain the differentiation of bacterial communities (Supplementary Data 6). Comparatively, the bacterial communities were much better grouped by the taxon of phytoplankton, which reflected the great impact of host-bacteria interaction as the most critical interspecific interaction in the phycosphere. Numerous studies based on *in situ* samples also revealed phytoplankton-species-dependent succession of bacterial communities^16, 19^. By performing RF, LEfSe and ROC on the big data, which was a widely recognized analytical pathway, we pinpointed 23 and 8 characteristic genera for dinoflagellate and diatom, respectively. Robust co-existences of some characteristic genera with their corresponding host phytoplankton can be verified by previous studies, some of which have further revealed adaptation mechanisms. *Polaribacter* is an example of, one of the most abundant characteristic genera for diatom, exhibiting high specificity for the binding and utilization of the diatom polysaccharide^20^. It dominates in *in situ* diatom blooms^21^ and becomes suppressed in dinoflagellate blooms^22, 23^, confirming its preference in the phycosphere of diatoms. On the contrary, phytoplankton associated *Roseibium* and *Balneola* are exclusively found in the phycosphere of dinoflagellates, with the presumed ability to regulate the synthesis of toxins in dinoflagellates^24, 25^. Characteristic genera did not attract much attention previously; most of them are less abundant in the phycosphere, some unknown, but interesting interactions with phytoplankton having potential significant ecological impacts are starting to reverse this.

Although the host-bacteria interaction was generally a greater influencer to bacterial communities in the phycosphere, the degree of influence varied with the host phytoplankton. First, independent analysis on the correlation between sampling locations and structural compositions of bacterial communities for either the dinoflagellate or diatom groups indicated that diatom-associated bacterial communities were more influenced by geographical differences. All the samples in the diatom group were taken from coastal waters of China, including ES (*n*=39), YS (*n*=14) and SC (*n* =57). Alpha-diversity showed significant differences between the bacterial communities in the ES and SS groups, and beta-diversity further indicated that all these three bacterial communities were structured differently (Supplementary Fig. 6, 7). In contrast, dinoflagellate-associated bacterial communities showed no significant pattern of variation with sampling locations, despite the wider range of geographic areas that samples were acquired (Supplementary Fig. 8, 9). Furthermore, characteristic genera in the dinoflagellate group were three times more than that of the diatom group and nearly one-third of these dinoflagellate-affiliated genera (7/23) did not belong to the three major groups of “archetypal phytoplankton-associated bacterial taxa” identified (Alpha-proteobacteria, Gamma-proteobacteria, Flavobacteriia). As for why the selection force of dinoflagellate was greater than that of diatom, the highly complex lifestyles of dinoflagellate could be a potential cause: dinoflagellates are highly motile^26, 27^; a large fraction of dinoflagellates are mixotrophs and can undergo phagocytosis to feed on microbes^28, 29^; many dinoflagellates also produce various toxins to influence the growth of other marine organisms^30, 31^. It suggests that metabolic specialists with specific capabilities, such as chemotaxis, adhesion, resistance to phagocytosis (capsule) or tolerance to toxins, are more likely to persistently exist in the phycosphere of dinoflagellates^32–35^. Most diatoms, however, are nonmotile and exclusively phototrophic. Only a small fraction of diatoms produce toxins and most diatom-derived organic matters are highly degradable by bacteria^36^. The frustule and transparent exopolymer particles of diatoms also provide a good habitat for bacterial colonization^37^. Therefore, metabolic generalists, which adapt to various environments, would dominate the phycosphere of diatoms.

Despite the great influence of host phytoplankton, many studies still disagree with the species-specific composition of bacterial communities in the phycosphere^38, 39^, which are reasonable since some conserved bacteria coexist with phytoplankton regardless of the taxon of phytoplankton and geographical location of sampling. However, most studies concerning core microbiota in the phycosphere were based on small datasets with certain similarities, which lacked universality. Our approach fully considers host-specificity and conservation of phytoplankton-associated bacterial communities whilst adopting all potential parameters concerning ecological importance, including abundance, frequency and connectivity. In addition, to verify the universal significance of these core genera, we proposed another exhaustive approach based on ubiquity-abundance (Ub-Ab) plot, global co-occurrence network and attribution of Bray-Curtis similarity to reveal their occupancy in bacterial community, connectivity to other genera and impact to community stabilization. The double detection approaches ensures that the 14 core genera we identified are solid. Inspecting current research on the 14 core genera reveal some interesting patterns. On one hand, all 14 core genera belong to the three major groups of “archetypal phytoplankton-associated bacterial taxa”, indicating that this study was not a subversion of the original knowledge, but a refinement of them. On the other hand, most of the core genera were rarely studied before. As of 12/20/2022, using the 14 genus names (Crocinitomicaceae as a substitute for *JN874375_g*) together with “phytoplankton” or “microalgae” as the keywords, literature retrieval on Web of Science (https://www.webofscience.com/wos) reveals that some genera have received greater attention, such as *Alteromonas* that has been reported for 118 times in the phycosphere, while there are nine genera studied for even less than one-tenth of the times of *Alteromonas*. Several neglected core genera showed salient ecological features, including *Marivita*, *Shimia* and *JN874375_g*. Some previous studies speculated that *Marivita* might play an important role in algal blooms^40^, but its metabolic and ecological functions in the phycosphere were virtually unknown except for providing vitamins^41, 42^ for phytoplankton. The knowledge on *Shimia* and *JN874375_g* was even more lacking. *Shimia* was only confirmed for its coexistence with phytoplankton^39^, and nothing was known about *JN874375_g*. None of the five core genera belonging to Flavobacteriia, which have been recognized as major groups in the phycosphere, were well studied. Similarly, *Pseudoruegeria* and *Planktotalea*, showed relatively high ecological features on connectivity with other genera and impact to community stabilization, but not occupancy in bacterial community. In addition, most Flavobacteria showed low culturabilities^43, 44^. Considering the great ecological significances of these core genera, research directions should be adjusted accordingly to shed more light on the previously neglected core genera.

The core genera were further categorized by the ecological features. Importantly, the classification approach based on big data analysis of community composition can accurately reveal three Tiers that were taxonomically different. Core genera belonging to Alteromonadaceae were all Tier-1 members, while core genera belonging to Flavobacteriaceae and Rhodobacteraceae distributed in all three tiers but were concentrated in Tier-2 and Tier-3, respectively. The great differences between bacterial taxon of the three tiers strongly suggest significant differences in metabolic capabilities. Digging into the current literatures concerning the 14 core genera, some intriguing patterns surfaced. Expect for *JN874375_g* as a rarely studied Flavobacteriia, all the remaining Tier-1 members are flagellated and therefore highly motile^45, 46^. In comparison, all Tier-2 members lack flagellum but showed gliding or wobbling movements^47, 48^, while all Tier-3 members are non-motile^49^. Since the Tiers were sorted by the significance of ecological features, it suggests motility may greatly contribute to earning ecological niches in the phycosphere. Meanwhile, significant overlaps of metabolic properties between Tiers were observed. Members of both Tier-1 and Tier-3 prefer phytoplankton-derived low-molecule-wight organics, such as DMSP^50–52^. All Tier-3 members and several Tier-1 members, such as *Alteromonas* and *Marinobacter*, have been reported for the abilities of inhibiting and promoting the growth of phytoplankton^4, 53, 54^, suggesting close relationships with phytoplankton. Similar lifestyle can be also interpreted from the higher niche overlap between Tier-1 and Tier-3. Comparatively, Tier-2 members, consisting of mostly Flavobacteria, had strong capabilities of degrading phytoplankton-derived high-molecule-weight organics but showed no capability of regulating the growth of phytoplankton^55, 56^. The relatively weaker connection with phytoplankton explains why Tier-2 is less homogenously selected and less overlapped with Tier-1. A possible cause for the more overlapped niches of Tier-2 and Tier-3 is that most genera belonging to these two Tiers are reported to be capable of interfering with the quorum sensing of other bacteria^57^, resulting in direct antagonistic effects. In summary, our study demonstrated that the ecological features of bacteria were strongly connected with their taxa and metabolic properties.

The identification of the core and characteristic microbiotas greatly expands our understanding of “archetypal phytoplankton-associated bacterial taxa”. The great difference in definition leads them to have quite different ecological niches. The core microbiota had wider ecological niche than the characteristic microbiotas, in both MA and PA samples. This confirmed the roles of core microbes as metabolic generalists that adapted to various environment, while defining characteristic microbes as metabolic specialists that gained advantages in the phycosphere of specific phytoplankton. The more general metabolic capability of the core microbiota led to the higher conservation and less volatility in the phycosphere, which could be confirmed by the lower mean absolute change rates of the total abundance of core microbiota in the PA samples arranged in time series (119% for core microbiota vs. 266% for diatom-affiliated characteristic microbiota vs. 196% for dinoflagellate-affiliated characteristic microbiota) (Supplementary Fig. 10). Therefore, although all these phytoplankton-associated bacterial taxa can be ideal targets in studying phytoplankton-bacteria interactions due to their ecological significance, tendency to choose between the core and characteristic genera should vary with specific research scenarios. For example, to compare the bacterial communities in the bloom seawaters to those in the non-bloom seawater, core genera would be more representative, while the characteristic genera are more suitable biomarkers for tracking the bacterial community succession in response to the variation of dominant phytoplankton during blooms. Collectively, the redefinition of “archetypal phytoplankton-associated bacterial taxa” lays solid foundation for studying the molecular and ecological mechanisms of phytoplankton-bacteria interactions and opens infinite possibilities for re-examining the significance of phytoplankton-associated bacteria from a global perspective.

## Methods

### Microalgal strains

In this study, 110 diatom strains affiliated to 28 genera and 86 dinoflagellate strains affiliated to 25 genera were isolated from the *in situ* seawater samples following the Daste’s method with some modification^58^: First, one drop of the *in situ* seawater sample was placed in the centre of a slide and surrounded by 4-5 drops of sterile F/2 medium^59^(F/2-Si medium for dinoflagellate strains). Then, under a light microscope, a single cell of microalgae was extracted from the *in situ* seawater droplet using a micropipette and transferred into a clean F/2 droplet. Next, the microalgal cell was washed by repeatedly pipetting from one F/2 droplet to another clean F/2 droplet 4-5 times to ensure the removal of free-living and loosely attached bacteria. Finally, the washed microalgal cell was transferred to one mL of sterile F/2 medium in a 24-well plate and incubated at 20 °C under a light intensity of 200 µmol photons m^-2^ s^-1^ with 12 h: 12 h light-dark cycles for 3-5 days before being semi-continuously cultured in a conical flask containing 30 mL of sterile F/2 medium under the same conditions. The *in situ* seawater samples for the isolation of diatom strains were taken from coastal waters of China, including the YS, ES and SS, while dinoflagellate strains originated from the Indian, Pacific and North Atlantic Oceans (Fig. 1a). Details are listed in Supplementary Data1. Since only bacteria tightly attached to the microalgal cells would be retained after repetitive washing during the isolation of microalgal strains, these bacterial communities were referred to as microalgae-attached (short as MA) in the following analysis.

### DNA Extraction and Sequencing

The MA bacterial communities were collected by filtering the microalgal cultures in exponential phases through 0.22 µm membrane filters. The filters were extracted for total genomic DNA by using a PowerSoil Kit (MoBio Laboratories, Solana Beach, CA, United States), strictly following the manufacturer’s instruction. High-quality DNA of the V4-V5 region of the 16S rRNA gene was amplified by PCR using the universal primers 515F (5’-GTGYCAGCMGCCGCGGTAA-3’) and 907R (5’-CCGTCAATTCMTTTRAGT −3’), and sequenced by BGI (The Beijing Genomics Institute, Shenzhen, Guangdong, China) via pair-end Illumina MiSeq platform. The raw sequencing data have been deposited in NCBI SRA under Bioproject PRJNA922461.

### Bioinformatic analyses of 16S rRNA gene amplicons

Raw sequencing data were processed using the QIIME2^60^software package (2021.2 release). Forward and reverse read pairs were spliced and the merged reads were quality filtered, denoised and clustered into ASVs at 99% similarities using the DADA2 algorithm^61^. Chimeras were identified and removed only when the parent sequences were at least two times as abundant as the corresponding chimeras. Taxonomic annotations for ASVs were conducted in two ways. The first method was based on a customized naive-Bayes classifier trained on 16S rRNA gene with the SILVA132 database^62^ release and trimmed to a length of 400 bp. The second method manually annotated all feature sequences on https://www.EzBioCloud.net/ using internal scripts from https://github.com/1996xjm/ezbiocloud and 97% similarity was set as the cutoff to identify the same genus. The ASVs that were not classified as bacteria, including those annotated as archaea, mitochondria, chloroplasts, and even without assigned domain, were removed. The phylogenetic metrics were constructed by aligning representative sequences using MAFFT^63^ and a phylogenetic tree was generated with FastTree^64^ after the multiple sequence alignment was masked and filtered.

### Alpha and Beta diversity analysis

Since the feature dataset for all samples varied between 145,136 and 8,387 reads, they were rarefied to 13,871 reads per sample for alpha- and beta-diversity analysis. Four different alpha-diversity indices (chao1, observed_otus, simpson, shannon) were calculated using the “core diversity metrics” function in QIIME2 and principal coordinate analysis (PCoA) was constructed by calculating the Unweighted-UniFrac distance matrix. Two-sided Wilcoxon rank-sum tests and PERMANOVA (permutation test with pseudo-Fratios) were used to compare the significance of differences in alpha- and beta-diversity indices between the dinoflagellate and diatom groups using the packages “vegan”, “reshape2” and “ggplot2” from R language. For every ANOVA, F values and degrees of freedom was provided in Supplementary Fig. 6.

### Network construction

The co-occurrence networks of ASVs in the dinoflagellate (*n*=86) and diatom (*n*=110) groups were constructed separately. Low-abundance ASVs with less than 10 reads were removed to reduce the complexity of the dataset. The initial networks were constructed using CoNet analysis combining Pearson and Spearman correlation with Bray-Curtis and Kullback-Leibler dissimilarity using the “CoNet” plugin in Cytoscape^65^ v3.9.0. The initial number of edges was set at 200 and the correlation threshold was automatically selected. Final networks were then obtained after filtering out invalid edges by performing a random permutation test (*p* value=0.05) and bootstrap score (*p* value=0.01) on the initial network. The parameters describing network topological characteristics included node number (the number of ASVs), edge number (the number of connections among all of the nodes), average degree (the average of the degrees of all nodes), clustering coefficient (the cohesiveness of networks), diameter (the maximum distance of any node), radius(the smallest non-zero value of all shortest paths), centralization (the tendency to have only one hub), average number of neighbors, heterogeneity(the tendency to have multiple hubs), characteristic path length(average shortest path of all nodes). Network nodes are clustered at the genus level for visualization.

### Identification of the characteristic genera for each phytoplankton group

To identify the characteristic genera with strong preferences for either dinoflagellates or diatoms, the difference in relative abundance for each genus in the two groups was determined by LEfSe using the OmicStudio tools at https://www.omicstudio.cn/tool/. LDA scores were generated by ranking the difference indices of differential abundance taxa to determine the effect sizes of these taxa and thereby distinguish between groups. Wilcoxon test was used to identify the taxa with significantly different relative abundances between the two groups. Genera with a *p* value < 0.05 by Wilcon rank sum test and LDA score > 3 were treated as the distinguished genera. Meanwhile, the random forest model with R package “randomForest” was used to generate a decision tree algorithm to generate MDG for each variable, ranking each genus from the highest to lowest. The tenfold cross verification with five repeats was then performed; according to the cross-validation curve and the principle of parsimony, the top 50 distinguished genera were selected to minimize cross-validation error and were identified as the characteristic genera. The visualization of variance values and relative abundance for each character genus were finished at https://www.chiplot.online/. Finally, the distinguished genera obtained in both analyses were determined as the characteristic genera. To verify whether the characteristic genera had ecologically meaningful preferences for dinoflagellates or diatoms, the ROC curve analysis was performed by “pROC” package in R. If the entire area under the ROC curve (AUC) including the confidence interval (95%) was greater than 0.5, the genus would be solidly confirmed as a characteristic genus. Meanwhile, the closer AUC is to 1, the stronger preference of the genus is for the specific host phytoplankton.

### Identification of the core genera

The EIGs were first determined based on the abundance, frequency, and connectivity in the bacterial communities of either dinoflagellate or diatom groups. First, the average relative abundance of each genus in the dinoflagellates (*n*=86) and diatoms (*n*=110) group was calculated, and genera with average relative abundance (ARA) greater than 1% were identified as the HAGs. Next, the frequency of each genus was calculated based on its occurrence or non-occurrence in all samples within the two groups. Genera with frequencies greater than 50% were regarded as the HFGs. The average relative abundance and frequency of each genus were calculated and visualized by the “ggplot2” and “microeco” package in R. Finally, all nodes of ASVs in the co-occurrence networks were clustered at the genus level and genera with degree ⩾6 were identified as network hubs and were referred to as the HCGs^66^. The EIGs of the dinoflagellate and diatom groups, respectively, were identified by taking the collections of the HAGs, HFGs and HCGs of corresponding groups. The core genera were determined by taking the intersection of EIGs of both groups.

### Qualification and classification of the global features of the core genera

Three global features (occupancy in bacterial community, connectivity to other genera and impact on community stabilization) of the core genera were characterized based on all the MA samples and each core genus was classified according to the level of each feature. First, the Ub-Ab curves were used to illustrate the relationships between ubiquity and abundance of the 14 core genera, as described by Li et al^67^. In the Ub-Ab curves, two pairs of abundance versus ubiquity cutoffs (0.01% | 40% and 1% | 15%) were used to determine the genera with low abundance but high ubiquity, and those with high abundance but low ubiquity. Genera that satisfied two, one and none of the cutoff pairs were defined as high-, medium-, and low-occupancy genera, respectively. Secondly, a global co-occurrence network was constructed on the genus level (same method as above), whilst clustering and visualization were performed based on the interrelationship types (copresence or mutual exclusion) between different genera. In the global co-occurrence network, high-, medium-, and low-connectivity genera were revealed based on the node degree cutoffs of 6 and 3. Finally, the contribution of each genus to the Bray-Curtis similarity of bacterial communities was calculated based on the method from Shade^68^ with some modifications. Different from the original method, we did not pre-order the genera just based on their frequencies in the samples. The first genus in the plot was determined by individually calculating the Bray-Curtis similarity of each genus and picking out the largest contributor. Then, the contribution of the remaining genus, together with the first genus, to the community stability was calculated by dividing the Bray-Curtis similarity of these two genera with the Bray-Curtis similarity of the whole genera), and the largest contributor was determined as the second genus. The third to the last genus were also determined in this way, resulting in a smoothly flattened Bray-Curtis similarity contribution curve. In this Bray-Curtis similarity contribution curve, cutoffs of last increasement of 3% and 2% were used to distinguish high-, medium-, and low-impact genera.

### Exploration of the ecological niches of core genera

The ecological niches of the core genera were demonstrated in three aspects, including the niche width, the degree of niche overlap as well as the positioning illustrated by the composition of ecological processes for community assembly. The ecological niche width and the degree of niche overlap were calculated based on the ‘Levins’ niche width index, which was used to determine precisely the mechanisms used to allocate available resources to the community and to confirm whether different species assemblages used habitat resources evenly (or asymmetrically). The calculations were conducted with the R package “spaa”. Then, the composition of ecological processes for community assembly was assessed by a null modeling framework iCAMP^69^, based on phylogenetic bins. The relative abundance of bins controlled by each process was aggregated to assess its impact on the overall community assembly. Separate tables of different ASVs were used to calculate the ecological null model. Results of the same type were summarized and then compared between the different community assemblies.

### Analysis of the *in situ* bloom data

*In situ* bloom data were collected from continuous monitoring of bacterial communities at the ‘Kabeltonne’ station (54° 11’ 17.88’’ N, 7° 54’ 0’’ E) from 2010 to 2013 (Supplementary Data 2), which was carried out by Teeling^70^. Algal blooms which were dominated by diatoms, dinoflagellates, and brown algae occurred frequently, and varied with time. According to the authors’ description, prefiltered seawaters (10 μm) were first filtered through 3 μm membranes to collect particle-attached bacteria (short as PA, 137 samples), and then through 0.22 μm membranes to collect free-living bacteria (short as FL, 91 samples). Raw reads of 16S rRNA genes for these samples were downloaded from NCBI (BioProject ID PRJNA266669-PRJNA266671) as typical *in situ* bloom data and were re-analyzed following the above methods to verify the ecological significance of phytoplankton-associated bacterial taxa.

## Data availability

All data used in this analysis are publicly available. The raw sequencing data of 16S rRNA amplicons from the microbial communities of dinoflagellates and diatoms generated in this study have been deposited in NCBI SRA under Bioproject PRJNA922461. Detailed data information is provided in the Supplementary Data 1.

## Acknowledgment

Funding for this study was provided by the National Natural Science Foundation of China (92051118, 32070113), and Guangdong Science and Technology Department (2022B1515020017).

## Author Contributions

H.W. conceived the experiment; X.Y., J.X., Y.C. and G.C. performed the experiment and data analysis, H.G. provided most of the dinoflagellates strains, X.Y. and Y.C. wrote the draft of the manuscript, H.W., X.Y., Y.C. and all other authors revised the manuscript.

## Ethical statement

This article does not contain any studies with human and animals performed.

