## Supplementary material for "Redefinition of archetypal phytoplankton-associated bacteria taxa based on globally distributed dinoflagellates and diatoms": Supplemenatary figures

\# These two authors contributed equally.

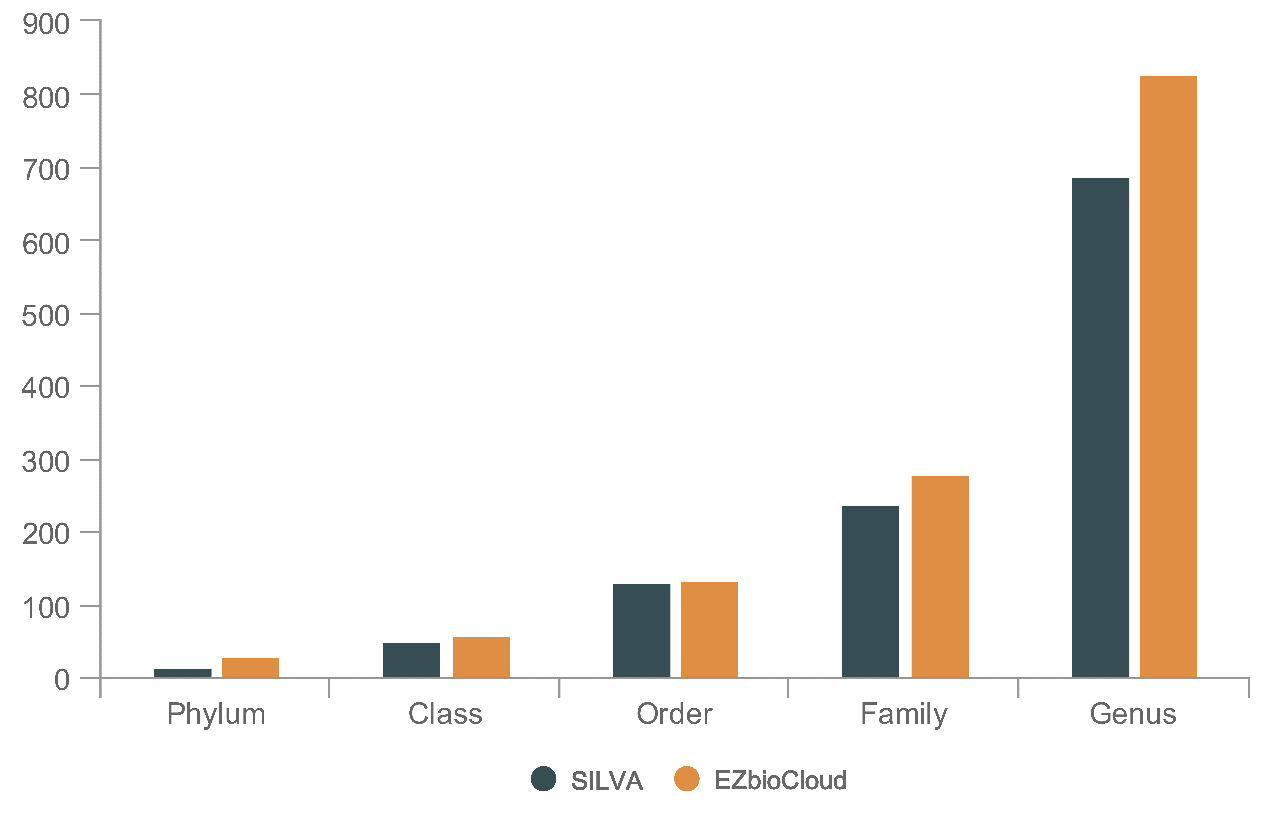

Supplementary Fig. 1. Comparison of annotation results of ASVs from 196 Microalgae-attached samples in two databases, SILVA and EZbioCloud.

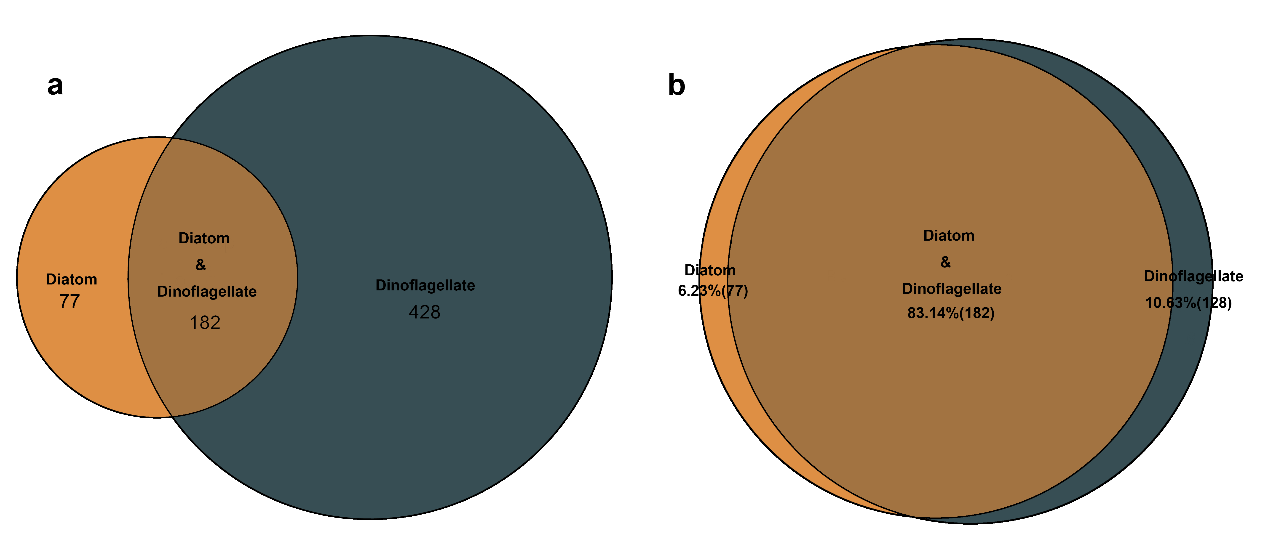

Supplementary Fig. 2. Comparison of the diversity of bacterial communities in the physcophere of dinoflagellates and diatoms. a, Venn diagram showing the numbers of genera in the dinoflagellate and diatom groups. b, Sequencing ratio of the same and different genera to total sequencing in the dinoflagellate and diatom groups.

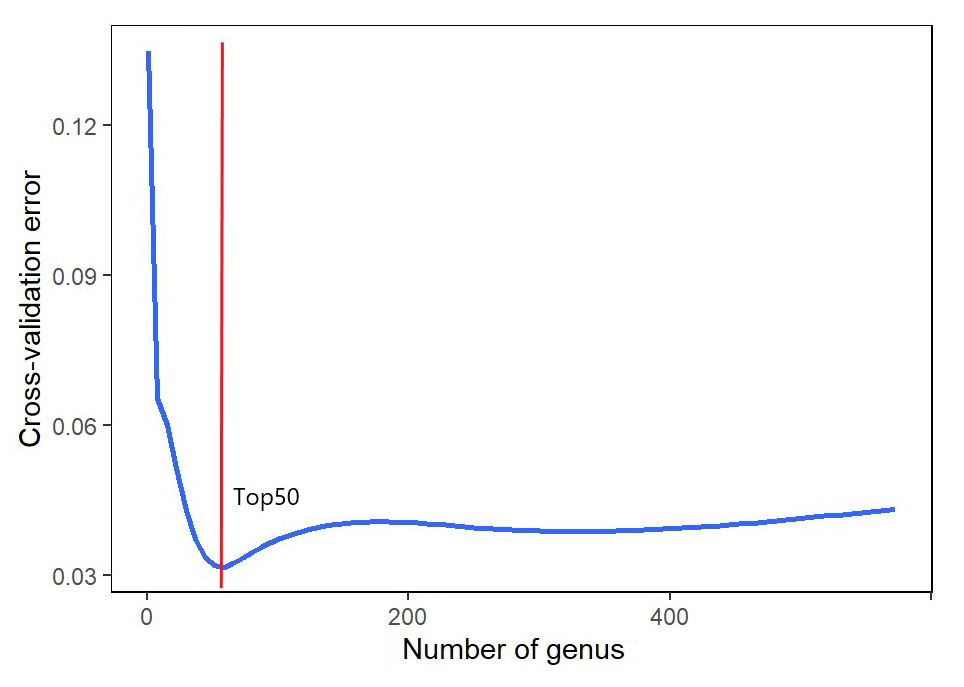

Supplementary Fig. 3. A ten-fold cross-validation curve with five repetitions based on the MeanDecreaseGini of each genus calculated in the random forest and sorted from highest to lowest. The top 50 genera were selected as representatives of the entire data set according to the horizontal coordinates corresponding to the lowest point of the curve (OOB estimate of error rate: 3.7%).

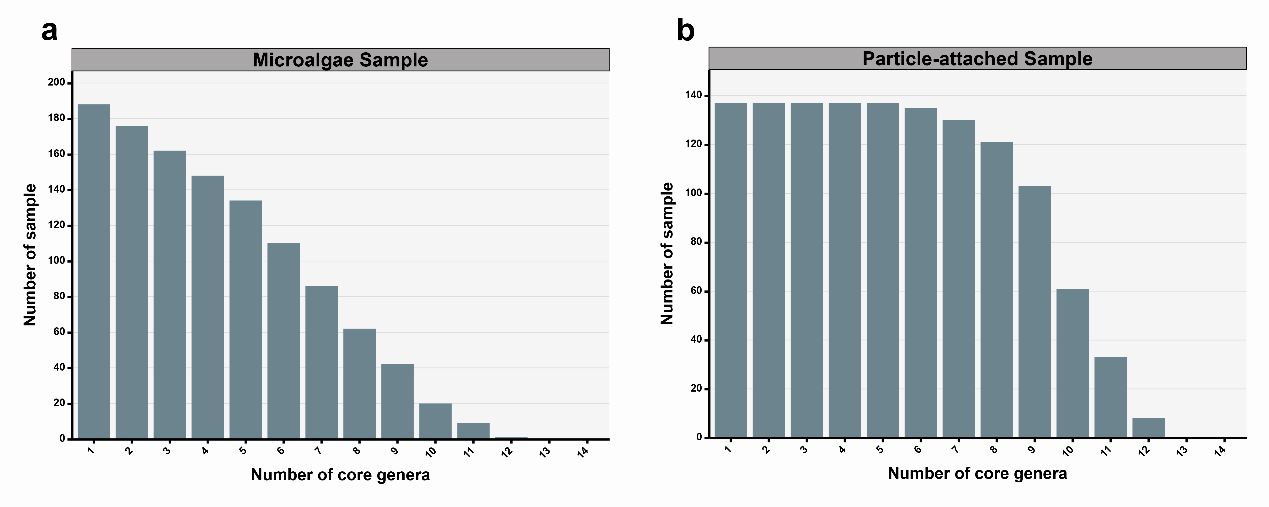

Supplementary Fig. 4. Distribution of core genera in the MA(a) and PA(b) samples.

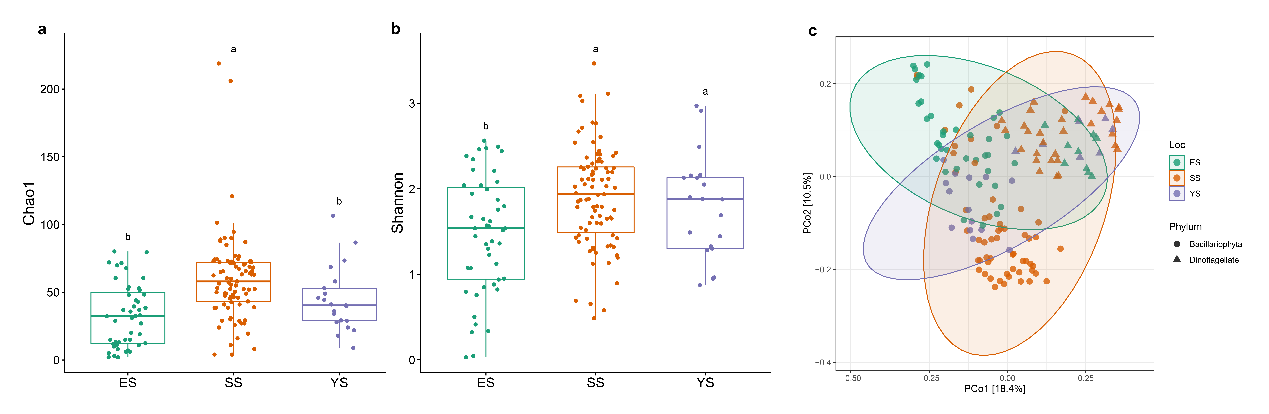

Supplementary Fig. 5. The α-diversity and β-diversity of bacterial communities in three sea area samples shared by dinoflagellates and diatoms samples. a, b represents the "Chao1" and "Shannon" indices, respectively, different letters indicate significant differences (*P* = 0.05, one-way ANOVA). c, d The Unwei- ght_Unifrac index plotted against the PCoA plot. PERMANOVA for significance of differences test.

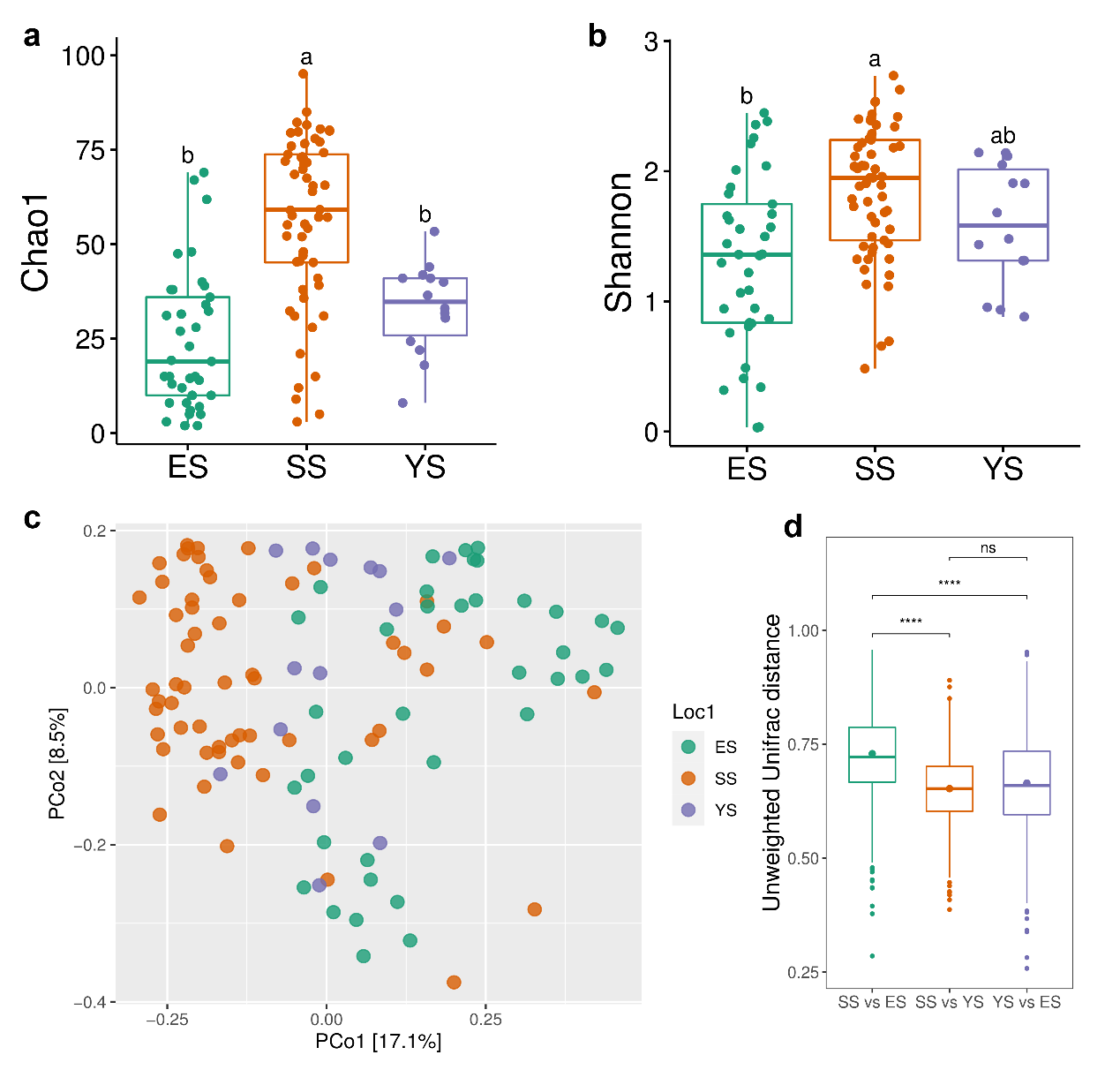

Supplementary Fig. 6. The α-diversity and β-diversity of bacterial communities in three different marine groups within the diatom group. a and b represent the "Chao1" and "Shannon" indices, respectively. Different letters indicate significant differences (*P* = 0.05, one-way ANOVA). c and d, the Unweight_Unifrac index plotted against the PCoA plot. PERMANOVA for significance of differences test. Colors represent different groups and stars represent significance levels. **** *P* ≤ 0.0001 *** *P* ≤ 0.001, ** *P* ≤ 0.01, * *P* ≤ 0.05, no star and transparent shading indicates *P* > 0.05 (not significant).

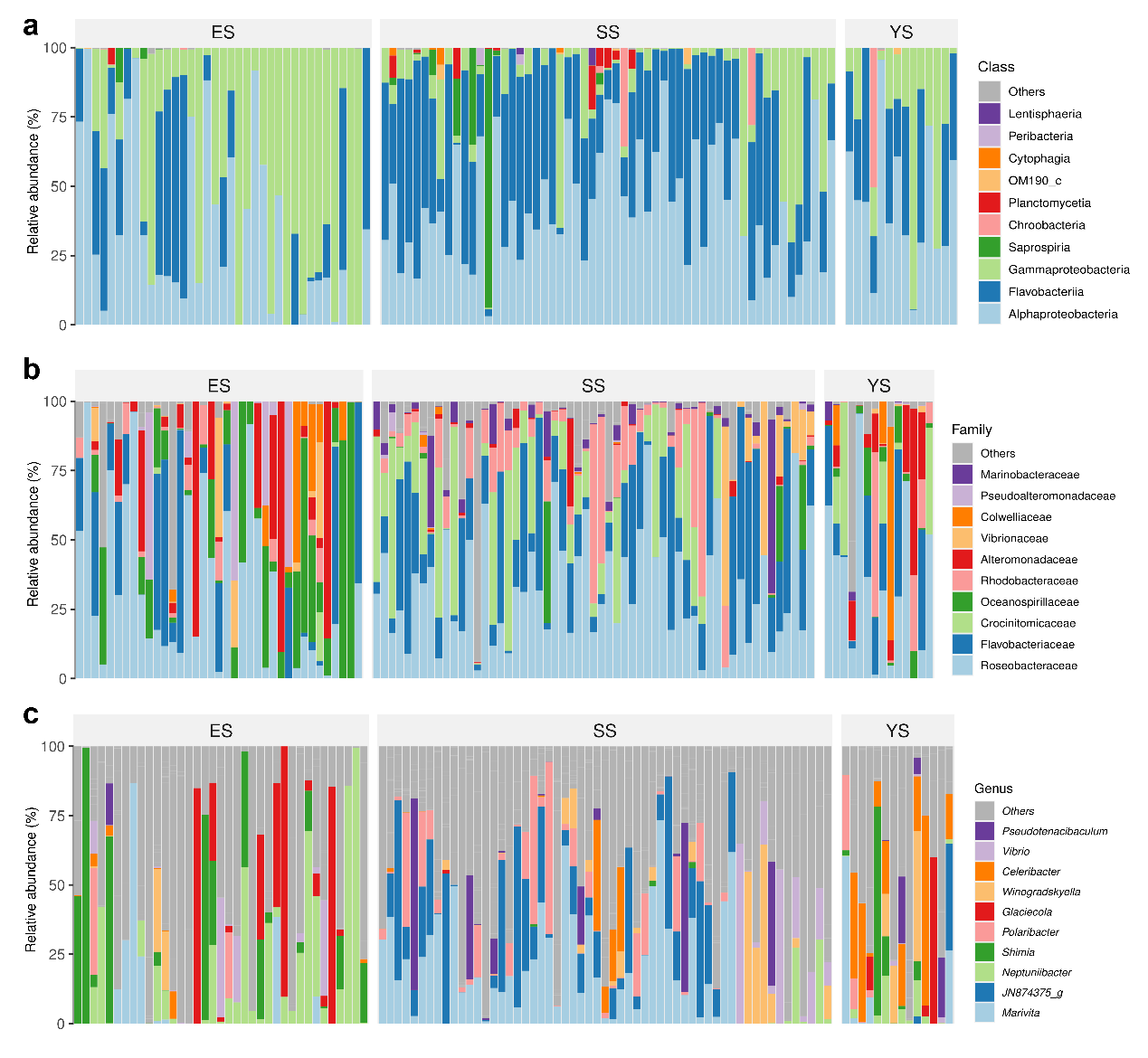

Supplementary Fig. 7 Histogram of species composition of the phycosphere for diatom samples grouped by sea area. a, b and c represent the top 10 bacterial group at the order, family, and genus levels, respectively.

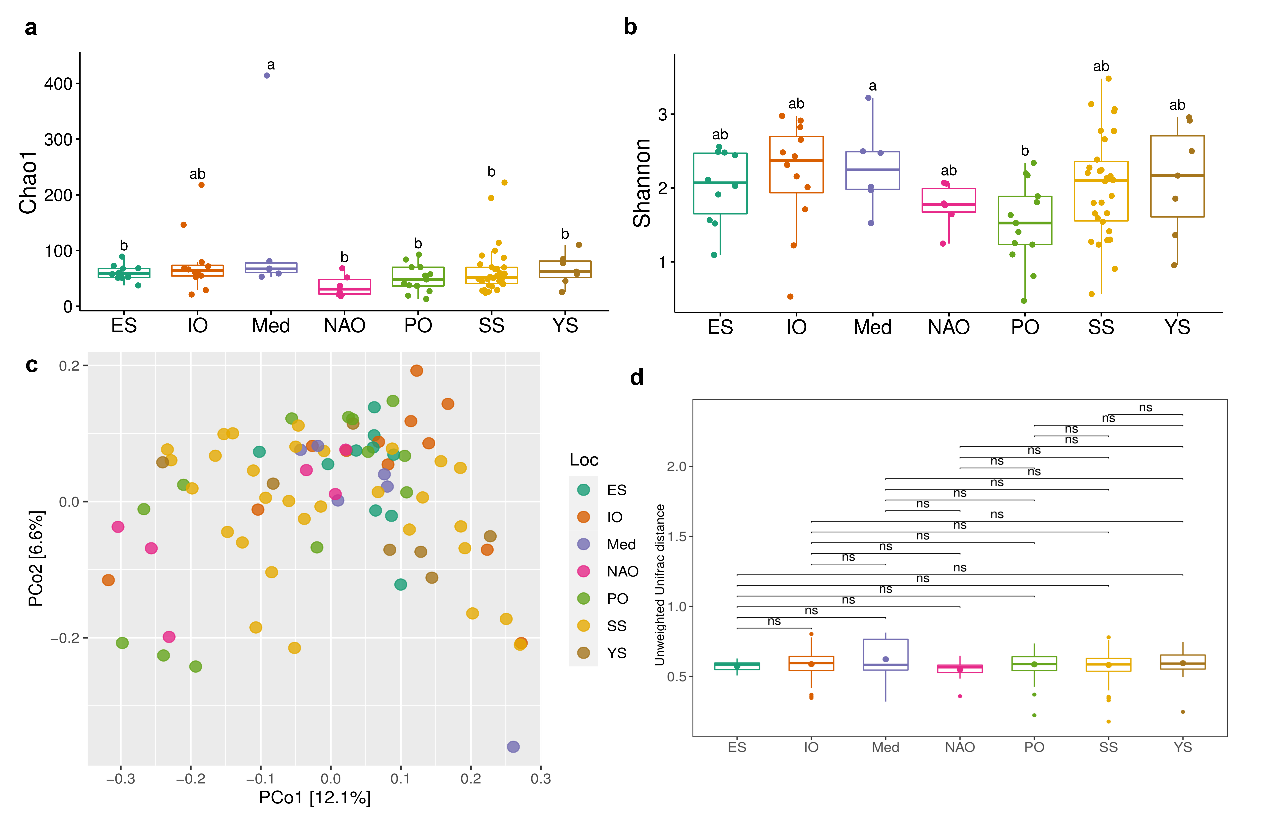

Supplementary Fig. 8 The α-diversity and β-diversity of bacterial communities in three different marine groups within the diatom group. a and b represent the "Chao1" and "Shannon" indices, respectively, different letters indicate significant differences (*P* = 0.05, one-way ANOVA). c and d, the Unweight_Unifrac index plotted against the PCoA plot. PERMANOVA for significance of differences test, Colors represent different groups and stars represent significance levels, **** *P* ≤ 0.0001 *** *P* ≤ 0.001, ** *P* ≤ 0.01, * *P* ≤ 0.05, no star and transparent shading indicates *P* > 0.05 (not significant).

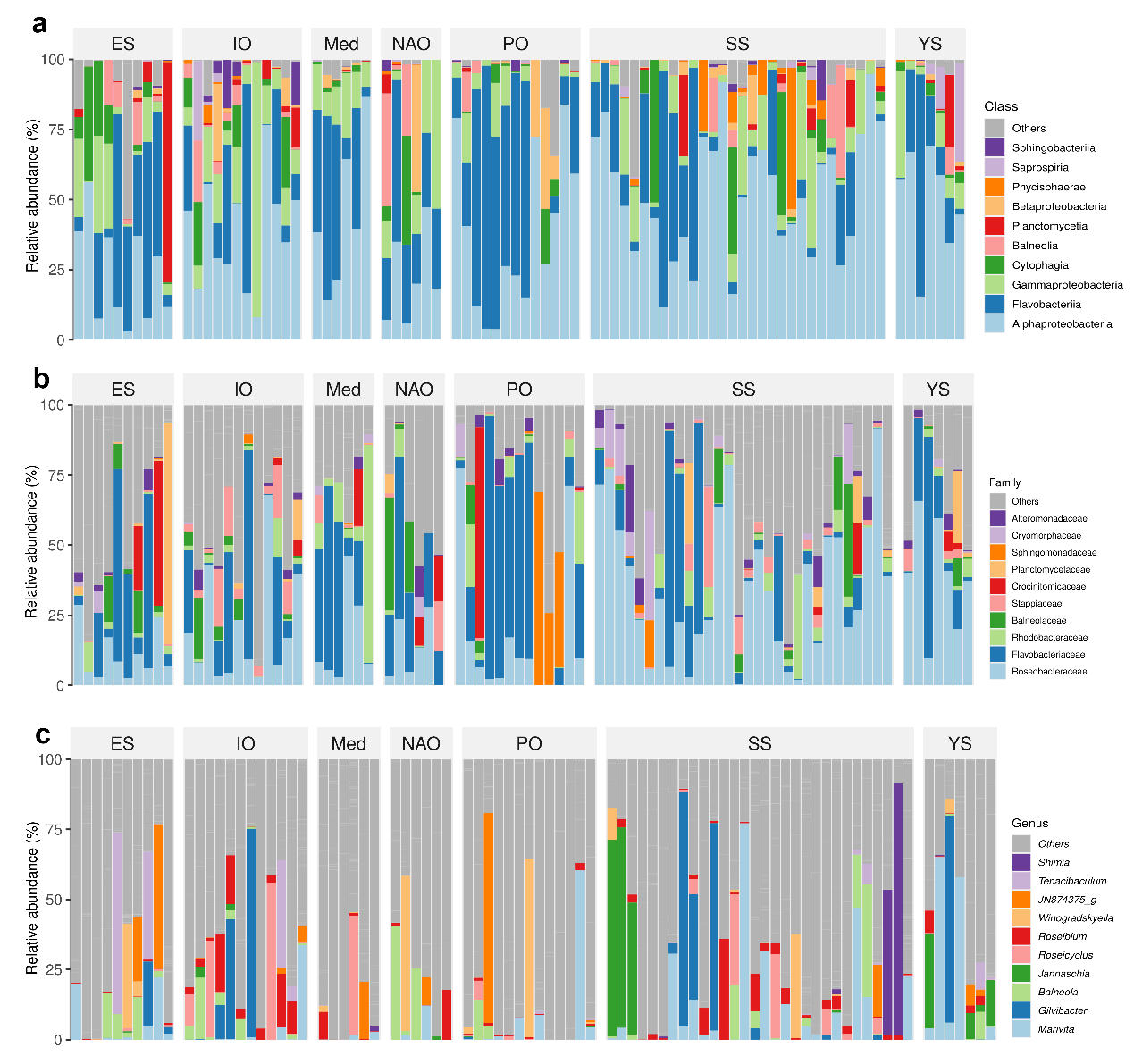

Supplementary Fig. 9 Histogram of species composition of the phycosphere for dinoflagellate samples grouped by sea area. a, b and c represent the top 10 at the order, family, and genus levels, respectively.

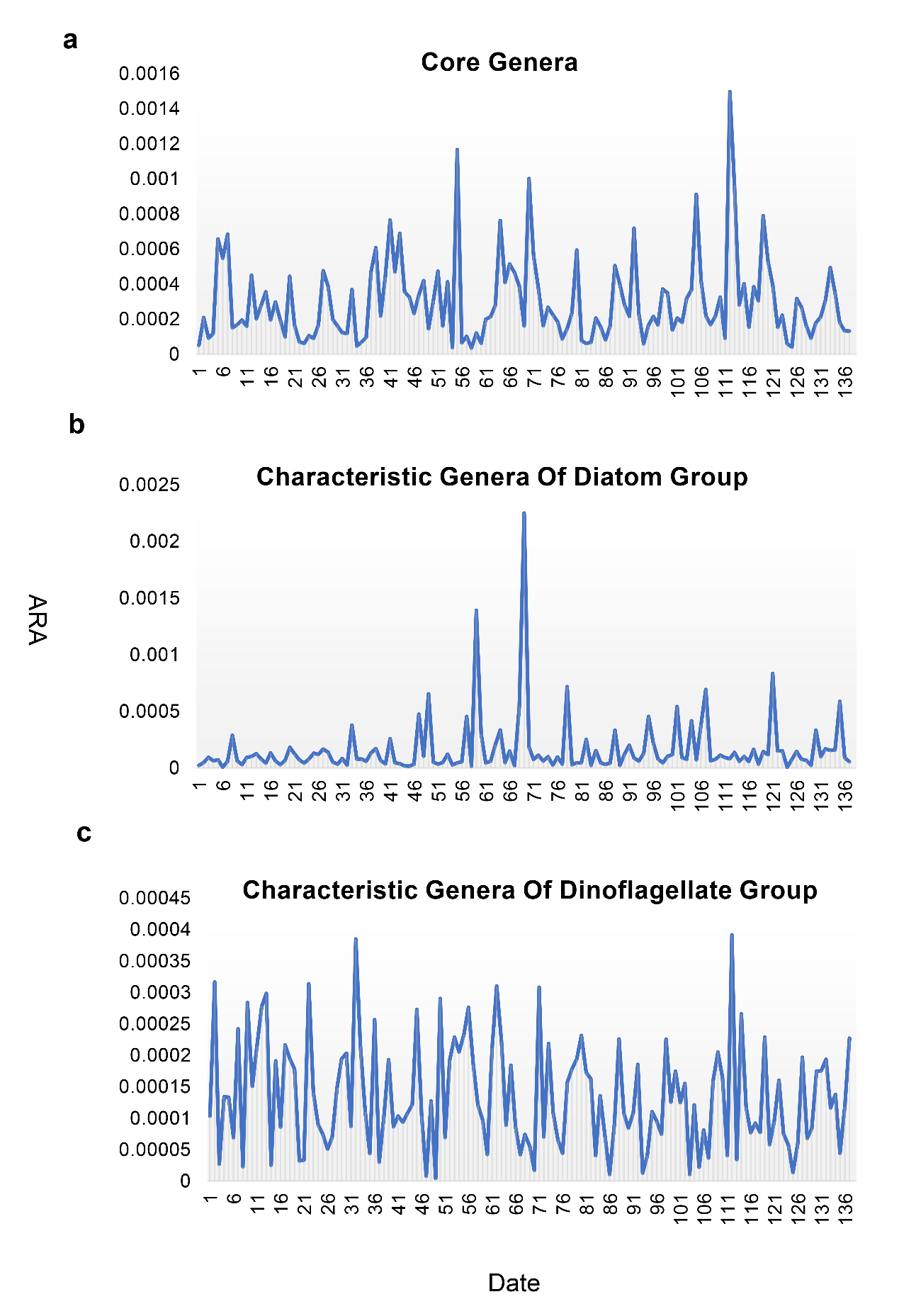

Supplementary Fig. 10 Variation curves of the mean relative abundance (ARA) of core genera (a) and characteristic genera (b, c) in PA samples of the time series.
